# Effects of MCHM on yeast metabolism

**DOI:** 10.1101/609800

**Authors:** Amaury Pupo, Kang Mo Ku, Jennifer E. G. Gallagher

## Abstract

On January 2014 approximately 10,000 gallons of crude 4-Methylcyclohexanemethanol (MCHM) and propylene glycol phenol ether (PPH) were accidentally released into the Elk River, West Virginia, contaminating the tap water of around 300,000 residents. Crude MCHM is an industrial chemical used as flotation reagent to clean coal. At the time of the spill, MCHM’s toxicological data were limited, an issue that have been addressed by different studies focused on understanding the immediate and long-term effects of MCHM on human health and the environment. Using *S. cerevisiae* as a model organism we study the effect of acute exposition to crude MCHM on metabolism. Yeasts were treated with MCHM 3.9 mM in YPD for 30 minutes. Polar and lipid metabolites were extracted from cells by a chloroform-methanol-water mixture. The extracts were then analyzed by direct injection ESI-MS and by GC-MS. The metabolomics analysis was complemented with flux balance analysis simulations done with genome-scale metabolic network models (GSMNM) of MCHM treated vs non-treated control. We integrated the effect of MCHM on yeast gene expression from RNA-Seq data within these GSMNM. 181 and 66 metabolites were identified by the ESI-MS and GC-MS procedures, respectively. From these 38 and 34 relevant metabolites were selected from ESI-MS and GC-MS respectively, for 72 unique compounds. MCHM induced amino acid accumulation, via its effects on amino acid metabolism, as well as a potential impairment of ribosome biogenesis. MCHM affects phospholipid biosynthesis and decrease the levels of ergosterol, with a potential impact in the biophysical properties of yeast cellular membranes. The FBA simulations were able to reproduce the deleterious effect of MCHM on cell’s growth and suggest that the effect of MCHM on ubiquinol:ferricytochrome c reductase reaction, caused by the under-expression of *CYT1* gene, could be the driven force behind the observed effect on yeast metabolism and growth.

## 1 Introduction

On January 2014 approximately 10,000 gallons of crude 4-Methylcyclohexanemethanol (MCHM) and propylene glycol phenol ether were accidentally released into the Elk River, West Virginia, contaminating the tap water of around 300,000 residents (Cooper, 2014)◻. Crude MCHM is an industrial chemical used as flotation reagent to clean coal (Christie et al., 1989)◻. More than 300 people in the affected area visited emergency departments with reports of symptoms potentially related to the spill, including mild skin, gastrointestinal and respiratory symptoms that were resolved with no or minimal treatment (Thomasson et al., 2017)◻. At the time of the spill, MCHM’s toxicological data were limited, an issue that have been addressed by different studies focused on understanding the immediate and long-term effects of MCHM on human health and the environment (Weidhaas et al., 2016)◻.

MCHM is considered a moderate-to-strong dermal irritant, causes fetal malformations in rats when orally exposed to 400 mg/kg/day (“Eastman Crude MCHM Studies,” 1990). The highest concentration of MCHM detected in tap water was 0.0294 mM (Whelton et al., 2015)◻. Crude MCHM is not a dermal irritant to humans at the concentrations in the water reported after the spill (Monnot et al., 2017)◻. In the evaluation of different cell lines, HEK-293, HepG2, H9c2 and GT1-7 only the highest dose of MCHM (1 mM) elicited a statistically significant decrease in cell viability, when compared to the control (1% DMSO) (Han et al., 2017)◻. MCHM induced DNA damage-related biomarkers in human A549 cells, indicating that it is related to genotoxicity (Lan et al., 2015)◻. MCHM affected larval visual motor response in an acute developmental toxicity assay with zebrafish embryos (Horzmann et al., 2017)◻. MCHM mainly induced chemical stress related to transmembrane transport activity and oxidative stress in yeast (Lan et al., 2015)◻.

The budding yeast *Saccharomyces cerevisiae* is one of the most intensively investigated, well-consolidated and widely used eukaryotic model organism. Its use has allowed the gain of insights in basic cellular mechanisms such as cell cycle progression, DNA replication, vesicular trafficking, protein turnover, longevity and cell death (Denoth Lippuner et al., 2014)◻ or even more complex process like neurodegenaretive disorders (Fruhmann et al., 2017). Being among the first components of the biota to be exposed to environmental pollutants, bacteria and fungi are common model organisms for eco-toxicological assessments (Braconi et al., 2016)◻. A number of features makes *S. cerevisiae* an ideal model for functional toxicological studies, such as: being unicellular, the ease of genetic manipulation, availability of a huge repertoire of dedicated experimental tools, protocols, software and databases, a high degree of functional conservation with more complex eukaryotes, among others (Braconi et al., 2016)◻. The effect of tens of pesticides have been studied in *S. cerevisiae* by a battery of omics approaches, including transcriptomics, chemogenomics, proteomics and metabolomics (reviewed in (Braconi et al., 2016)◻).

Focused on the analysis of the whole repertoires of endogenous or exogenous metabolites that are present in a biological system at a given time point metabolomics serves as a link between genotype and phenotype (Aliferis and Chrysayi-Tokousbalides, 2011; Ibáñez et al., 2013). Metabolomics is an extremely useful tool in the analysis of the metabolic modifications induced by potentially toxic compounds (Nicholson et al., 2002)◻. These studies include the effect different fungicides (Queiroz et al., 2012), Cu^2+^ exposure (Farrés et al., 2016), tolerance to representative inhibitors (Wang et al., 2015)◻, ethanol tolerance (Kim et al., 2016; Ohta et al., 2016), among others.

Flux balance analysis (FBA) (Fell and Small, 1986; Savinell and Palsson, 1992; Varma et al., 1993; Varma and Palsson, 1993) with genome-scale metabolic network models (GSMNM) allows the simulation of the metabolism at a systemic level, for the understanding of diverse phenomena and making predictions (Yilmaz and Walhout, 2017)◻. There are more than twenty genome-scale metabolic network models reconstructed for *S. cerevisiae* to date (Heavner and Price, 2015)◻. The consensus yeast metabolic network stands out with 14 compartments, more than 3700 reactions, >2500 metabolites and >1100 genes (Heavner et al., 2013)◻.

The accuracy of FBA predictions can be improved by the integration of experimental data (Yilmaz and Walhout, 2017)◻. Several methods have been developed to this end, allowing the integration of transcriptomics data: such as E-Flux (Colijn et al., 2009)◻, omFBA (Guo and Feng, 2016)◻ and transcriptional regulated flux balance analysis (TRFBA) (Motamedian et al., 2017)◻; proteomics data: GECKO (a method that enhances a genome-scale metabolic models to account for enzymes as part of reactions) (Sánchez et al., 2017)◻; and metabolomics data: unsteady-state flux balance analysis (uFBA) (Bordbar et al., 2017)◻.

In the present work we study the effect of MCHM on metabolism using yeast as a model organism, combining metabolomics tools with FBA simulations on genome-scale metabolic network models of yeast constrained by RNA-Seq data.

## 2 Materials and Methods

### 2.1 MCHM treatment

Wildtype yeast from the S288c background (BY4741 strain *his3, ura3, leu2, met15*) (Brachmann et al., 1998)◻ were grown in YPD to exponential phase (OD 0.4 – 0.6) then treated with crude 4-Methylcyclohexanemethanol (crude MCHM provided directly from Eastman Chemical) 3.9 mM for 30 minutes or left untreated. Six independent biological replicates were done per treated and untreated group. After 30 minutes 5 optical units of cells were collected, washed with deionized water, flash frozen in liquid nitrogen and stored at −80⁰C for extraction within the next 24 hours.

### 2.2 Metabolites extraction

Lipid and polar metabolites were extracted with a 1:2:0.8 mixture of chloroform: MeOH: H2O, following a modified version of a published protocol (Bourque and Titorenko, 2009)◻. HPLC grade chloroform and methanol were from Sigma-Aldrich. All the steps were done using glassware, to avoid polymers contamination. The extractions were performed in 15 mL Kimble™ Kontes™ KIMAX™ Reusable High Strength Centrifuge Tubes from Fisher Scientific. Half of the original protocol volume values were used. For extractions headed to GC-MS analysis 50 μL of ribitol internal standard (10 mg/mL) were added. 3 mL of the polar and 3 mL of the lipid phase were collected per sample. The polar phase were dried in SpeedVac (ThermoFisher Scientific). The lipid phase was dried overnight in a fume hood. For ESI-MS experiments, but not for GC-MS, the dried polar phases were re-suspended in 500 μL of MeOH and the lipid phases were re-suspended in 500 μL 1:1 chloroform: MeOH. All extracts were stored at −20 °C for analysis within 48 hours.

### 2.3 ESI-MS

Samples were analyzed by direct injection of the resuspended extracts in a Thermo Fisher Scientific Q-Exactive, with an ESI (electrospray ion source), using positive and negative modes. For polar compounds in positive mode the injection speed was 10 μL/min, the scan range was 50 – 750 m/z, no fragmentation, 140,000 resolution, 1 microscan, AGC target 5*10^5^, maximum injection time of 100, sheath gas flow rate of 10, aux gas flow rate of 2, no sweep gas flow, spray voltage 3.60 kV, capillary temperature of 320°C, S-lens RF level 30.0. For polar compounds in negative mode most parameters remain the same, except for spray voltage: 3.20 kV, capillary temperature: 300 °C, S-lens RF level: 25.0. For lipid compounds in positive mode the following parameters were modified; scan range: 150.0 – 2,000.0 m/z, sheath gas flow rate: 15, aux gas flow rate: 11, spray voltage: 3.50, capillary temperature: 300°C, S-lens RF level: 25.0. For lipid compounds in negative mode the previous parameters were kept, except for the spray voltage, which was set to 3.20 kV.

50 scans were obtained per sample and later averaged with Thermo Scientific Xcalibur 2.1 SP1. Averaged spectra in positive and negative mode were processed for polar and lipid fractions, separately, with xcms 3.2.0 (Smith et al., 2006)◻. Peaks were identified within each spectrum using the *mass spec wavelet* method from the MassSpecWavelet 1.46.0 R package (Du et al., 2006)◻. Peaks were grouped with the *Mzclust* method, followed by *groupChromPeaks*. All features were plotted to be visually inspected. The intensity values of each feature in each sample was obtained with the *featureValue* method as the integrated signal area for each representative peak per sample. The feature intensity and feature definition tables were saved as CSV files. Features were identified via MetaboSearch 1.2 (Zhou et al., 2012)◻, with the list comprising the average mz values for each feature as a query, with 5 ppm of error, positive or negative mode and using the four online databases available as options in the program: HMDB, Metlin, MMCD and LipidMaps. After the feature identification, feature intensity tables (keeping only identified features) coming from the same biological replicate (both positive and negative modes from polar and lipid fractions) were merged as a single intensity table.

Six biological replicates per group for MCHM treated and untreated controls were used. The experiment were repeated twice with consistent results. These biological replicates are not the same used in GC-MS experiments.

### 2.4 GC-MS

50 μL of Methyl heptadecanoate 2 mg/mL was added as internal standard to each lipid sample before derivatization. Lipid and polar fractions were derivatized with BSTFA (DATTA et al., 2012)◻ and MSTFA (Xue et al., 2015)◻, respectively. For BSTFA derivatization dried extracts were treated with 200 μL N,O-bis(trimethylsilyl)trifluoroacetamide with 1% of trimethylchlorosilane at 75 °C for 30 min. For MSTFA derivatization dried extracts were treated with 50 μL methoxyamine hydrochloride (40 mg/ml in pyridine) for 90 min at 37 °C, then with 100 μL MSTFA + 1% TMCS at 50 °C for 20 min. Derivatized samples were analyzed using a GC-MS (Trace 1310 GC, Thermo Fisher Scientific, Waltham, MA, USA) coupled to a MS detector system (ISQ QD, Thermo Fisher Scientific, Waltham, MA, USA) and an autosampler (Triplus RSH, Thermo Fisher Scientific, Waltham, MA). A capillary column (Rxi-5Sil MS, Restek, Bellefonte, PA, USA; 30 m × 0.25 mm × 0.25 μm capillary column w/10 m Integra-Guard Column) was used to detect polar metabolites. For water-soluble metabolite analysis, after an initial temperature hold at 80 °C for 2 min, the oven temperature was increased to 330 °C at 15 °C min^−1^ and held for 5 min. For lipid-soluble metabolite analysis, after an initial temperature hold at 150 °C for 1 min, the oven temperature was increased to 320 °C at 12 °C min^−1^ and held for 7 min. Injector and detector temperatures were set at 250 °C and 250 °C, respectively. An aliquot of 1 μL was injected with the split ratio of 70:1. The helium carrier gas was kept at a constant flow rate of 1.2 mL min^−1^. The mass spectrometer was operated in positive electron impact mode (EI) at 70.0 eV ionization energy at m/z 40-500 scan range.

Peak identification and grouping, and feature intensities calculation were performed with Thermo Scientific™ Chromeleon™ (Version 7.2, Thermo Fisher Scientific, Waltham, MA, USA). Features were identified against a locally characterized set of central metabolites (targeted metabolomics), when possible. Other features were identified querying NIST database (untargeted metabolomics). Feature intensity tables were saved as CSV files, keeping only the identified features.

Features intensities from lipid and polar fractions were normalized against its corresponding internal standards (methyl heptadecanoate for lipid and ribitol for polar fractions) and then the ones coming from the same biological replicate (both lipid and polar fractions) were merged as a single intensity table.

Six biological replicates per group for MCHM treatment and untreated controls were used. The experiment was repeated three times with consistent results. These biological replicates are not the same used in ESI-MS experiments.

### 2.5 Metabolomics data processing

Feature intensity tables from ESI-MS and GC-MS were processed with MetaboAnalyst 4.0 (Chong et al., 2018)◻. Missing intensity values were replaced by half of the minimum positive value in the original data. Up to 5% of the features with near-constant intensity values among the samples were filtered out. Samples were normalized by sum and data scaled by Pareto scaling. Samples are compared by univariated analysis (t-test and fold change) and multivariate analysis: Partial Least Squares Discriminant Analysis (PLS-DA), Sparse Partial Least Squares –Discriminant Analysis (sPLS-DA), Empirical Bayesian Analysis of Microarray (EBAM), Random Forest classification and Significance Analysis of Microarray (SAM), as implemented in MetaboAnalyst 4.0. Relevant metabolites were chosen among those that changes significantly due to the treatment and the relevant metabolites from the multivariate analysis.

The Pathway Analysis were performed with MetaboAnalyst 4.0 using the name of the relevant compounds from the ESI-MS and GC-MS combined. The *Saccharomyces cerevisiae* pathway library was used, as well as the hypergeometric test for the over representation analysis and relative-betweenness centrality for the pathway topology analysis.

Some pathways are represented as Escher maps (King et al., 2015)◻ with the thick and color of the edges as a function of the respective MCHM treated vs untreated control flux ratio values.

### 2.6 Transcriptomics

A fraction of previously reported data was used, including only the samples with wildtype S288c (S96 *lys5)* cells in YPD treated or not with MCHM (Pupo et al., 2019)◻. The RNA-seq of S96 was carried out on hot phenol extracted RNA (Rong-Mullins et al., 2017)◻. The raw data is accessible at https://www.ncbi.nlm.nih.gov/geo/query/acc.cgi?acc=GSE108873, containing count data generated via Rsubread and the differential expression data generated via DESeq2. MA plot and KEGG Pathway Enrichment Analysis were done with R packages *ggpubr* and *clusterProfiler* (Yu et al., 2012)◻, respectively.

### 2.7 Flux Balance Anaylsis

For our FBA simulations we used the consensus genome-scale metabolic model of *Saccharomyces cerevisiae*, yeastGEM, version 8.3.0 (Kerkhoven et al., 2018)◻. The simulations were performed with the COBRApy python package (Ebrahim et al., 2013)◻, using yeastGEM definition of growth as the objective function to be maximized.

The upper bounds of reactions from yeastGEM were modified in correspondence with gene expression of related genes from our RNA-Seq data. For this integration of RNA-Seq and FBA we adapted the *E-Flux* method developed by *Colijn et al.* (Colijn et al., 2009)◻. Briefly, every reaction is associate with a set of genes which products (enzymes or transporters) make the reaction possible. In the simplest case only one gene or none at all are associated, meaning that the enzyme catalyzing the reaction is a single poly-peptide entity or that the reaction is spontaneous, respectively. When the enzymes are heteromeric the gene coding for the different subunits are associated by an “AND” keyword, and the maximum reaction flux is driven by the gene with the lowest expression of the set. When the reaction can be driven by more than one protein the corresponding gene (gene sets) are associated by the “OR” keyword, and the maximum reaction flux is a function of the sum of the corresponding gene (gene sets) expressions. If there is no expression value for a given gene the average expression of the corresponding experimental group is used instead.

The resulting upper reaction bounds were normalized between zero and 1000 (the default upper bound in the yeastGEM model). Two models came out as the result of this procedure, one for MCHM treated yeast and one for the untreated control.

Default solutions are determined for each model using the *optimize* method from COBRApy and with the default yeastGEM media. Phenotype phase plane of Growth vs D-Glucose exchange was calculated with the *production_envelope* method and the corresponding graphics generated with *ggpubr*.

Upper bounds of selected reactions were manually modified to test for the importance of such reactions in growth.

All fluxes are in *mmol/(gDW*\**hour)*.

## 3 Results

### 3.1 MCHM affects yeast metabolism

To assess how MCHM treatment affects metabolism, 181 and 66 metabolites were identified by the ESI-MS and GC-MS procedures, respectively. There is almost no overlap between both set of compounds, as only eight metabolites were detected by both procedures: adenosine, citric acid, L-proline, myristic, palmitic, palmitoleic and stearic acids and uridine. A total of 238 metabolites were consistently detected by our combined analysis, comprising 15 out of the 20 standard amino acids and 51 variants of phospholipids, among other lipid and polar compounds (Supplemental Table 1).

Features from the MS spectra were detected, grouped, identified and their intensities calculated as described in Materials and Methods. Intensities were normalized to facilitate multivariate analysis (see Materials and Methods, Supplemental Figure 1 and 2). The 25 metabolites that change most significantly due to MCHM treatment per experiment type, as detected in ESI-MS and GC-MS, are shown in Figure 1. Biological replicates per group are consistent, clustered in the heatmaps (Figure 1). The concentration of some metabolites are increased due to the MCHM treatment, including nine amino acids (N, V, A, D, T, G, Y, S, Q), TCA cycle intermediates (citric and malic acids), glutathione and inosine (Figure 1), while the levels of other are decreased, such as: phospholipids and phosphatidic acids, ergosterol, L-ornithine, urea, uracil and myo-inositol.

**Figure 1:**
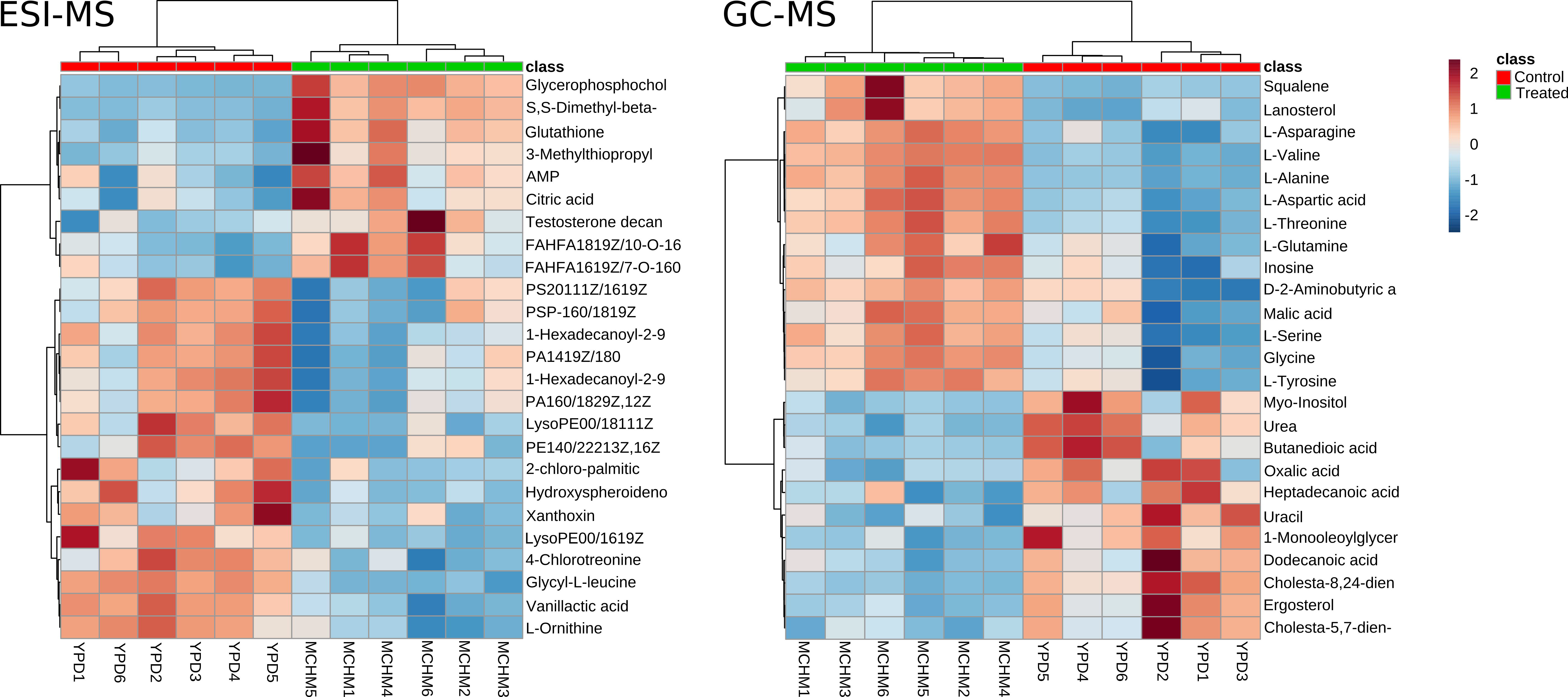
Heatmaps with normalized expression of the 25 most significant metabolites from ESI-MS and GC-MS experiments.

In order to assess the proper differentiation of the MCHM treated vs untreated control groups and to test the relevancy of significant metabolites by an alternative way we performed a Partial Least Squares Discriminant Analysis (PLS-DA, Figure 2). The MCHM treated and untreated control groups can be clearly separated in both ESI-MS and GC-MS experiments (Figure 2 A and B). This group separation was confirmed with Orthogonal-Orthogonal Projections to Latent Structures Discriminant Analysis (OPLS-DA, Supplemental Figure 3) and Sparse Partial Least Squares-Discriminant Analysis (sPLS-DA, Supplemental Figure 4 A, B), showing that such separation reflects the effect of MCHM treatment and is independent of specificities in the implementation of the Partial Least Squares Discriminant Analysis methodology. PLS-DA allows finding out the metabolites responsible for the separation of the groups. Compounds with VIP (variable importance in projection) scores for the first component greater than 1 were selected as relevant metabolites with PLS-DA (Figure 2 C and D). A total of 34 and 15 compounds were found relevant by PLS-DA in ESI-MS and GC-Ms experiments, respectively. From these some were not previously shown in the 25^th^ most significant (Figure 1) such as: 2-isopropylmalate and acetylcarnitine. The list of relevant metabolites was consolidated with the selected metabolites from sPLS-DA (Supplemental Figure 4 C and D), and the significant features identified by Random Forest (Supplemental Figure 5), Empirical Bayesian Analysis of Microarray (EBAM, Supplemental Figure 6) and Significance Analysis of Microarray (SAM, Supplemental Figure 7).

**Figure 2:**
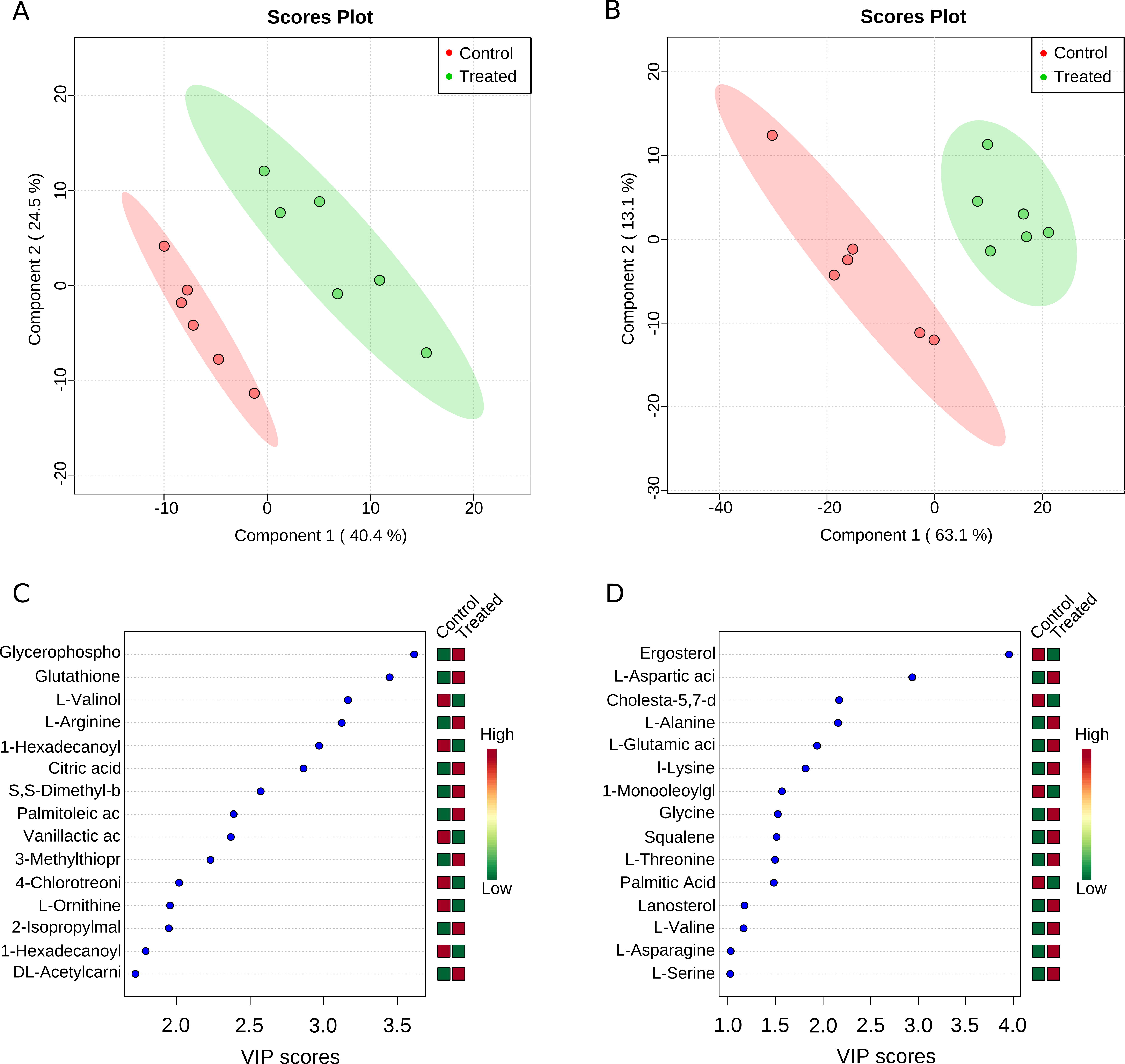
Partial Least Squares Discriminant Analysis (PLS-DA). Scores plot between the first two components for A) ESI-MS and B) GC-MS respectively, with the 95 % confidence area shown and the explained variance shown in brackets in the corresponding axis labels. Important features identified by PLS-DA in C) ESI-MS and D) GC-MS experiments. The colored boxes on the right indicate the relative concentrations of the corresponding metabolite in each group under study.

Metabolites being significant in most of the statistical analysis were selected as relevant metabolites, which concentrations change due to MCHM treatment (Figure 3). There are 38 and 34 relevant metabolites from ESI-MS and GC-MS respectively, for 72 unique compounds. From these the levels of 40 are reduced and the levels of 32 are increased due to MCHM treatment. In addition to the metabolites mentioned above, relevant metabolites include fatty acids, branched fatty acid esters, d-glucose, AMP, guanosine, four new amino acids (R, E, K, and P, for a total of 13 out of 20 standard amino acids), 3-methylthiopropyl acetate, pyroglutamic acid, lactic acid, among others.

**Figure 3:**
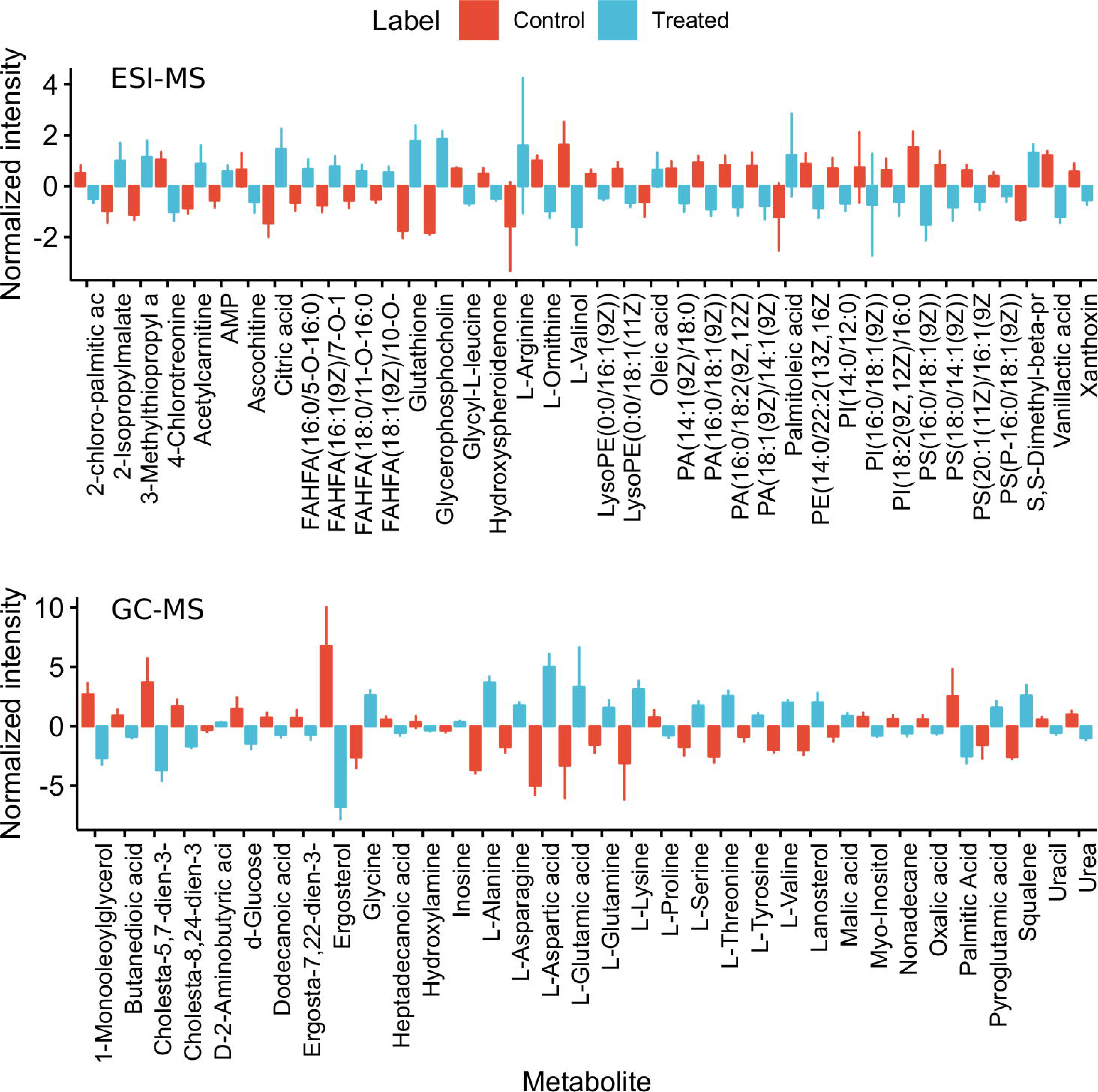
Normalized intensity of relevant metabolites from ESI-MS (upper panel) and GC-MS (lower panel).

These relevant metabolites were used as input for a pathway analysis (Figure 4), which combine pathway enrichment with pathway topology analysis. Six metabolic pathways are both statistically significant and with impact (Figure 4). Relevant amino acids dominated this analysis, as three of the pathways are the metabolism of amino acids (including A, D, E, R, P, G, S and T), the aminoacil t-RNA biosynthesis reactions has them as reactants, and the nitrogen metabolism has L-glutamate, L-glutamine and 2-oxoglutarate as intermediaries. The other relevant pathway is the glutathione metabolism, being the levels of glutathione, a critical redox agent, increased due to MCHM treatment.

**Figure 4:**
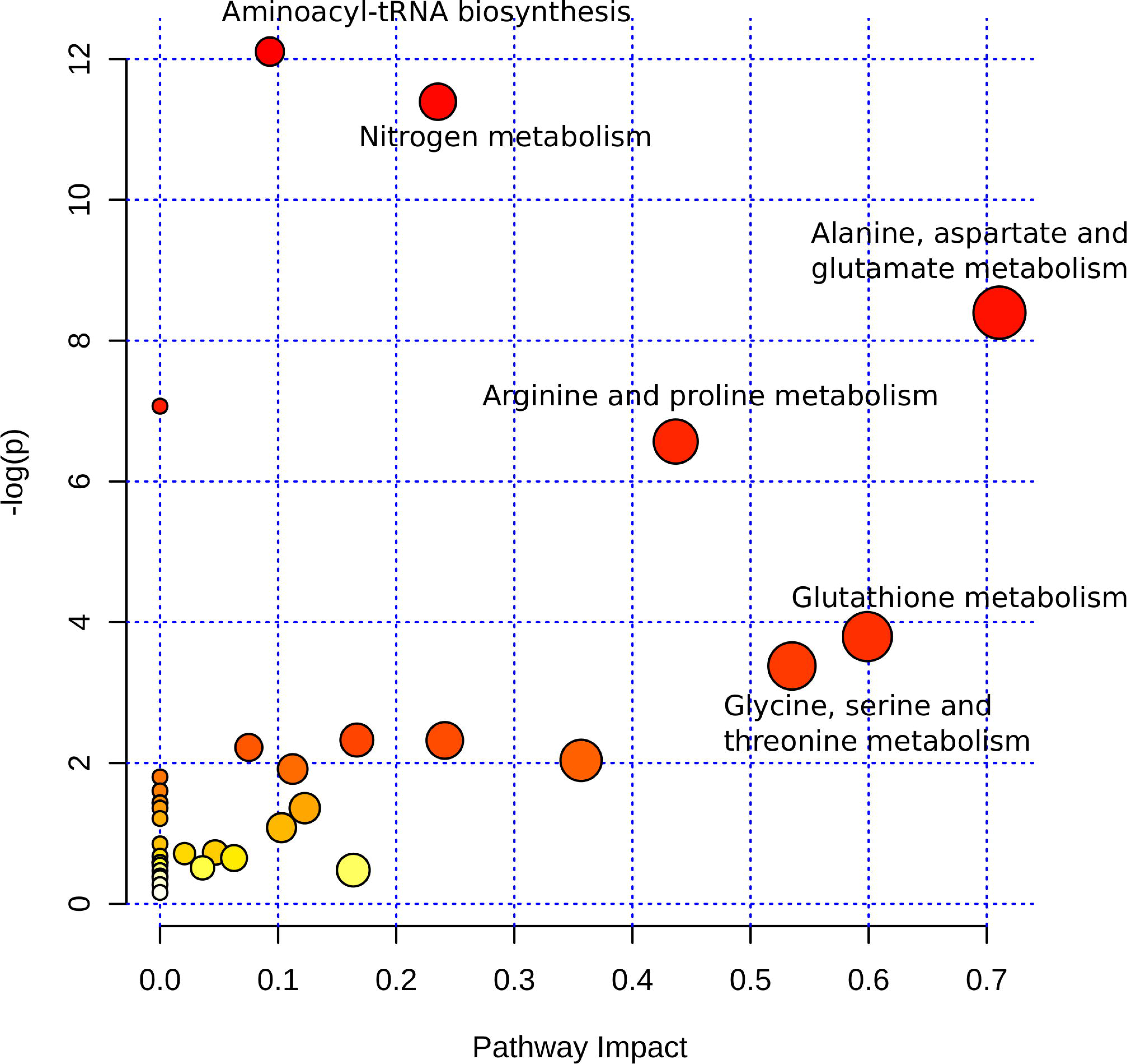
Pathway Analysis using relevant metabolites from ESI-MS and GC-MS combined. The six pathways with impact greater than zero and p < 0.05 are labeled.

### 3.2 Effect of MCHM on gene expression

We used a data set generated previously by our laboratory and available from https://www.ncbi.nlm.nih.gov/geo/query/acc.cgi?acc=GSE108873 (Pupo et al., 2019)◻. For this analysis we kept only the data regarding wild type S288c strain in YPD, treated or not with MCHM by 90 minutes.

From gene expression measurements for 3946 genes, 87 were up-regulated and 30 down-regulated due to MCHM treatment (Figure 5A, Supplemental Table 2) potentially affecting 18 metabolic pathways (Figure 5B). No pathway enrichment was found from the down-regulated genes.

**Figure 5:**
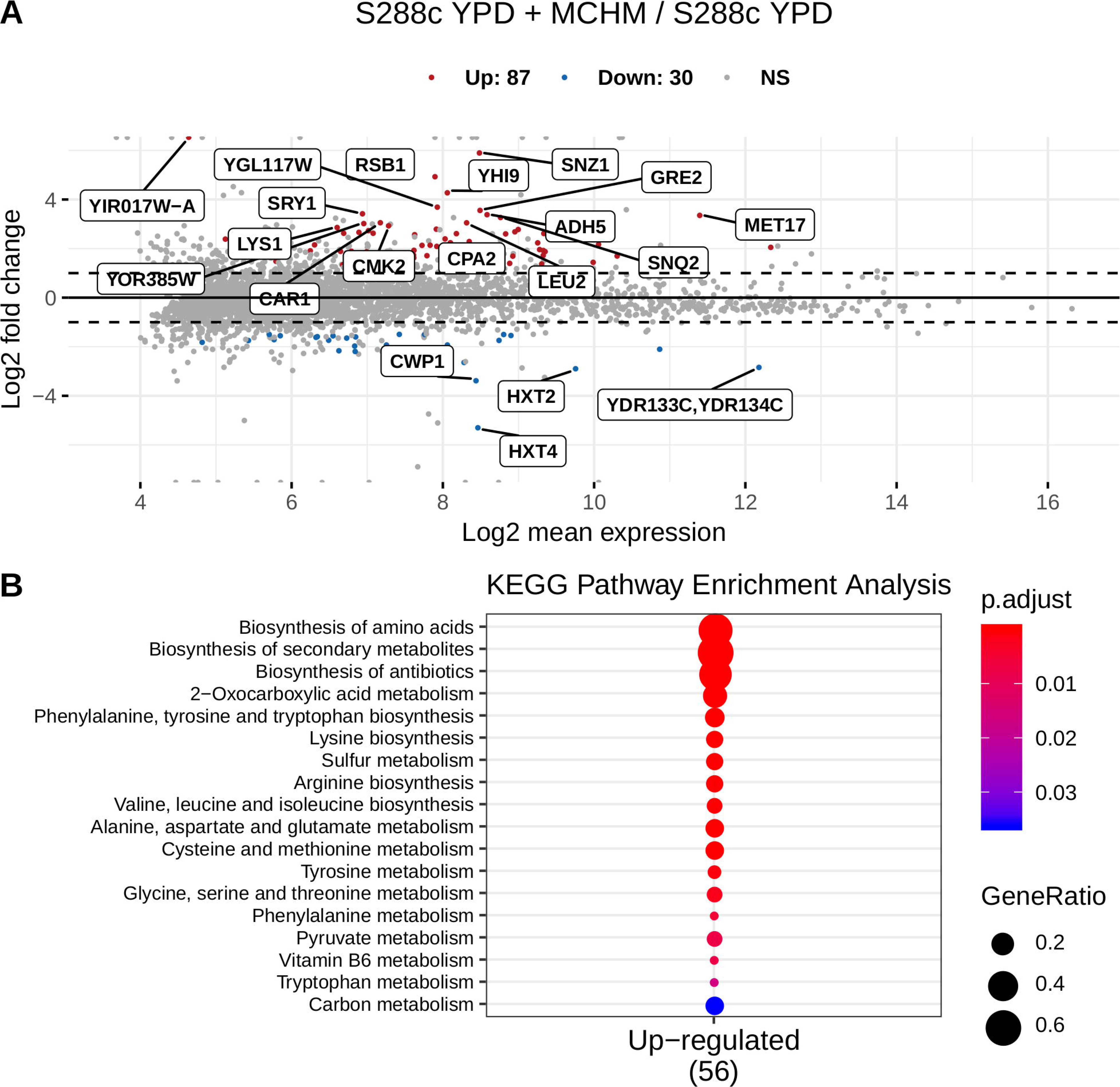
RNA-Seq gene expression data, represented as an MA plot (A). KEGG Pathway Enrichment Analysis for diferentially expressed genes (B). No enrichment was found for down-regulated genes.

Seven down-regulated genes are involved in ribosome biogenesis: *SDA1* and *RRP1*, involved in 60S ribosome biogenesis (Babbio et al., 2004; Horsey et al., 2004). ESF1, its depletion causes severely decreased 18S rRNA levels (Peng, 2004)◻. *BFR2*, involved in pre-18S rRNA processing and component of 90S preribosomes (Pérez-Fernández et al., 2007)◻. *MRD1*, required for production of 18S rRNA and small ribosomal subunit (Jin et al., 2002)◻. *NOP4*, constituent of 66S pre-ribosomal particles and critical for large ribosomal subunit biogenesis and processing and maturation of 27S pre-rRNA (Sun et al., 1994)◻. *NOP7*, component of several pre-ribosomal particles (Miles et al., 2005)◻. Loss of Sda1 function causes cells to arrest in G1 before Start and to remain uniformly as unbudded cells that do not increase significantly in size (Buscemi et al., 2000; Zimmerman and Kellogg, 2001)◻.

Among the rest of down-regulated genes there are two that codes for cell wall mannoproteins (*CWP1* and *TIR1*) and three involved in iron and zinc transport and homeostasis (*FTR1*, *ZRT1* and *IZH1*).

The up-regulated gene set is enriched in genes coding for enzymes of the amino acid biosynthesis pathways (28 out of 87) (Figure 5B, Supplemental Table 2): *ARG1*, *ARG5,6*, *ARG7*, *CPA1*, *CPA2*, *ASN1*, *GDH1*, *HIS4*, *HIS5*, *HOM2*, *HOM3*, *LEU1*, *LEU2*, *LEU4*, *LYS1*, *LYS2*, *LYS12*, *MET5*, *MET6*, *MET17*, *MET22*, *TRP2*, *TRP5*, *TMT1*, *ARO1*, *ARO3*, *ADE3* and *THR4*. These gene products participate in the biosynthesis of the amino acids: D, R, N, E, H, M, T, L, K, C, W, Y and F. Other three genes: *CAR1*, *MET3* and *MET14* are involved in R and M metabolism. *ARO8*, coding for the aromatic aminotransferase I, is also up-regulated and its expression is regulated by general control of amino acid biosynthesis (Iraqui et al., 1998)◻.

Five genes, which protein products abundance increases in response to DNA replication stress, are up-regulated due to MCHM treatment: *AHA1*, *ENO1*, *GRE2*, *PDR3* and *PDR16*. Other stress response related genes are also up-regulated: *CMK2*, *ICT1*, *TPO1*, *ENB1*, *SNQ2* and *QDR3*.

It is of note that genes coding for six mitochondrial enzymes (*MAE1*, *BAT1*, *ILV6*, *IDP1*, *GCV2* and *LYS12*) and three mitochondrial transporters (*GGC1*, *OAC1* and *ODC2*) are up-regulated. From these, MAE1 codes for the mitochondrial malic enzyme which catalyzes the decarboxylation of malate to pyruvate (in addition to its key role in sugar metabolism, pyruvate is a precursor for synthesis of several amino acids); *BAT1* and *ILV6* products are involved in branched-chain amino acid biosynthesis and *ODC2* codes the 2-oxodicarboxylate transporter, which exports 2-oxoglutarate and 2-oxoadipate from the mitochondrial matrix to the cytosol for use in glutamate biosynthesis and in lysine metabolism.

### 3.3 Modeling MCHM effect on yeast metabolism by Flux Balance Analysis

Using the expression data and the gene rules from the yeastGEM model (version 8.3.0) upper bounds were calculated for 2504 reactions of the model. Two new metabolic models were created from the original yeastGEM model, named *control* and *treated*, with the upper bounds of their reaction fluxes calculated from the corresponding gene expression data (as explained in Materials and Methods), and using as the objective function the maximization of growth. A summary of the result of FBA simulations with these models are shown in Tables 1 and 2. All the input and output fluxes are shown, with the involved metabolites, the calculated flux rates and their ranges. Flux ranges were calculated by Flux Variability Analysis with a fraction to optimum of 1. The objective function flux is shown. The growth was predicted to decrease due to the MCHM treatment, from a flux of 0.0704 to 0.0591 mmol/(gDW*hour) (Tables 1 and 2). The flux ratio of growth between treated and control was ∼0.839. So, MCHM treatment decreased yeast growth, consistent with the experimental results (Pupo et al., 2019)◻.

**Table 1:** FAB solution for the control model. [e] indicates extracellular compartment. All fluxes are in mmol/(gDW*hour).

| IN FLUXES |  |  | OUT FLUXES |  |  | OBJECTIVES |  |
| --- | --- | --- | --- | --- | --- | --- | --- |
| Name | Flux | Range | Name | Flux | Range | Name | Flux |
| oxygen [e] | 1.91 | [1.91, 1.91] | H2O [e] | 2.82 | [2.23, 2.82] | growth | **0.0704** |
| phosphate [e] | 1.2 | [0.0178, 1.2] | formate [e] | 1.74 | [1.74, 1.74] |  |  |
| D-glucose [e] | 1 | [1, 1] | carbon dioxide [e] | 1.35 | [1.35, 1.35] |  |  |
| ammonium | 0.388 | [0.388, 0.388] | diphosphate | 0.59 | [0, 0.59] |  |  |
| [e] |  |  | [e] |  |  |  |  |
| sulphate [e] | 0.00538 | 0.00538, 0.00538] | H+ [e] | 0.302 | [0.302, 1.06] |  |  |
|  |  |  | ethanol [e] | 0.17 | [0.17, 0.17] |  |  |

**Table 2:** FAB solution for the treated model. [e] indicates extracellular compartment. All fluxes are in mmol/(gDW*hour).

| IN FLUXES |  |  | OUT FLUXES |  |  | OBJECTIVES |  |
| --- | --- | --- | --- | --- | --- | --- | --- |
| Name | Flux | Range | Name | Flux | Range | Name | Flux |
| oxygen [e] | 1.4 | [1.4, 1.4] | H2O [e] | 1.93 | [1.77, 1.93] | growth | **0.0591** |
| D-glucose [e] | 1 | [1, 1] | carbon dioxide [e] | 1.45 | [1.45, 1.45] |  |  |
| phosphate [e] | 0.345 | [0.015, 0.345] | formate [e] | 1.25 | [1.25, 1.25] |  |  |
| ammonium [e] | 0.325 | [0.325, 0.325] | ethanol [e] | 0.57 | [0.57, 0.57] |  |  |
| sulphate [e] | 0.00451 | [0.00451, 0.00451] | H+ [e] | 0.212 | [0.212, 1.38] |  |  |
|  |  |  | diphosphate [e] | 0.165 | [0, 0.165] |  |  |

Our FBA simulations predict that the effect of MCHM on growth is diminished when the concentration of D-Glucose in the medium is decreased (Figure 6). There is a level of D-Glucose in the medium (∼0.5 mmol/(gDW*hour)) from which the growth of the MCHM treated and control models are the same.

**Figure 6:**
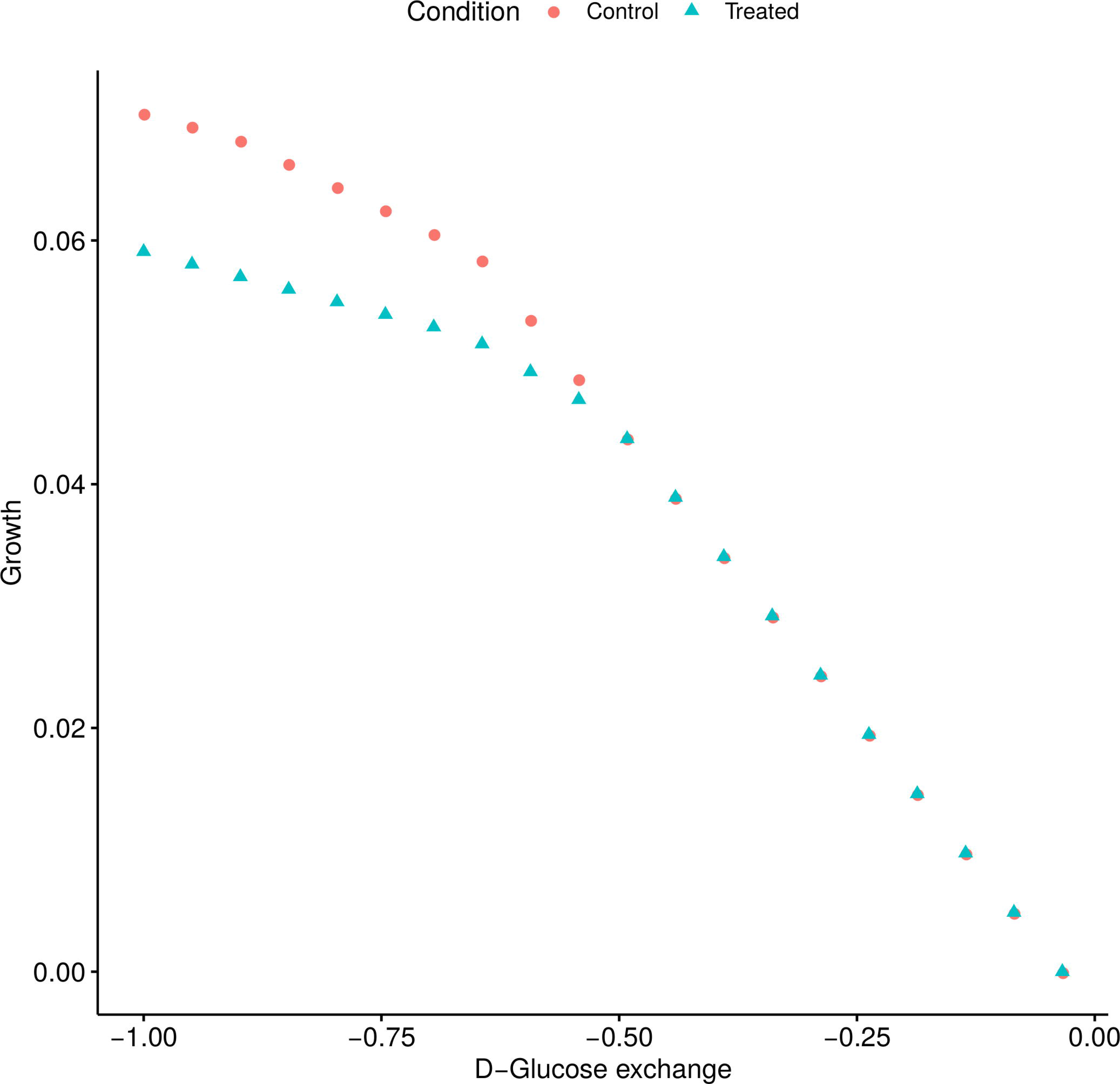
The effect of MCHM on yeast growth is predicted to depend on the concentration of D-Glucose in the medium. Phenotype phase plane of Growth vs D-Glucose exchange, from the FBA simulations with the control and treated models.

We then focused in the six significant pathways from the pathway analysis (Figure 4), to analyze the flux ratios between the FBA solutions of the treated vs the control models. The Escher maps representations (King et al., 2015)◻ of alanine, aspartate and glutamate metabolism, glutathione metabolism, glycine, serine and threonine metabolism, arginine and proline metabolism, nitrogen metabolism and the aminoacil t-RNA biosynthesis are shown in Supplemental Figures 8 – 13. As in any metabolic map the nodes are the metabolites and the edges connecting them are the reactions, with arrow heads indicating the reaction direction and labeled by the corresponding enzyme or transporter. The ratio of the fluxes passing throughout the respective reactions in the MCHM treated vs untreated control models are shown next to the enzyme names, and the color and width of the edges are scaled in function of such ratio values. All the relevant pathways have fluxes affected due to the treatment, fluxes that involved some relevant metabolites from the metabolomics studies. Only two reactions of glutathione metabolism are relevant in the solutions of these FBA simulations, both involved in the inter-conversion between glutathione and glutathione disulfide (Supplemental Figure 9). No flux goes through the other reactions in the pathway. A similar situation is present in nitrogen metabolism pathway, with flux passing only through glutamine synthetase and bicarbonate formation reactions (Supplemental Figure 12). In the rest of the analyzed pathways most of the reactions are active (with non-zero net fluxes) (Supplemental Figure 8, 10, 11 and 13). The fluxes of most reactions are decreased in the MCHM treated model vs the control (flux ratios < 1). There are many reactions which flux ratio (treatment/control) is the same ratio of the growth, the value 0.839. The extreme case is Aminoacil t-RNA biosynthesis (Supplemental Figure 13), where all the reactions have this flux ratio. These reactions having the same treated/control flux ratio as the treated/control growth ratio indicates that they are linked to the growth but does not ensure that any of these reactions is actually limiting it.

#### 3.3.1 Limiting reaction in FBA models

The reaction or reactions limiting the growth (limiting reactions) must be operating at the maximum allowed flux (upper bound value, calculated in function of the related gene expression levels) in the *treated* model. Two reactions operated at max flux in the model of the treatment (Table 3, last two rows). One of these, the ubiquinol:ferricytochrome c reductase, was also operating almost at maximum flux in the control model (Table 3, data row 3), and it is then the primary candidate to be the limiting reaction in our FBA simulations. Ubiquinol:ferricytochrome c reductase is part of oxidative phosphorylation pathway and contributes to the proton gradient formation through the mitochondrial membrane.

**Table 3:** Reactions operating within 0.1 units of the maximum allowed flux. Compartments legend: c, cytoplasm; e, extracellular; m, mitochondria. All fluxes are in mmol/(gDW*hour).

| Id | Name | Flux | Upper bound | Compartment | Model |
| --- | --- | --- | --- | --- | --- |
| r_0226 | ATP synthase | 4.375 | 4.375 | m, c | Control |
| r_0438 | ferrocytochrome-c:oxygen oxidoreductase | 6.930 | 6.930 | m, c | Control |
| r_0439 | ubiquinol:ferricytochrome c reductase | 3.465 | 3.536 | m, c | Control |
| r_0501 | glycine cleavage system | 0.449 | 0.449 | m | Control |
| r_0506 | glycine-cleavage complex (lipoylprotein) | 0.423 | 0.449 | m | Control |
| r_0507 | glycine-cleavage complex (lipoylprotein) | 0.423 | 0.449 | m | Control |
| r_0508 | glycine-cleavage complex (lipoylprotein) | 0.423 | 0.449 | m | Control |
| r_0773 | NADH:ubiquinone oxidoreductase | 0.730 | 0.730 | m | Control |
| r_1250 | putrescine excretion | 0.539 | 0.539 | e, c | Control |
| r_0439 | ubiquinol:ferricytochrome c reductase | 2.582 | 2.582 | m, c | Treated |
| r_0569 | inorganic diphosphatase | 0.330 | 0.330 | m | Treated |

To test if *ubiquinol:ferricytochrome c reductase* was the limiting reaction we modified its upper bound in the control model to the one it has in the treated (Table 4, third data row vs first and second data row). The growth rate decreased from 0.0704 to 0.0597, which is practically the same growth of the treated model, 0.0591. As can be seen modifying the maximum allowed flux of this reaction alone is enough to mimic the effect of the treatment in the growth, **confirming that ubiquinol:ferricytochrome c reductase is the limiting reaction** in our FBA simulations. We tried to recover the control phenotype (growth of 0.0704, Table 4, first data row) by setting the *ubiquinol:ferricytochrome c reductase* upper bound in the treated model to the one it has in the control one (Table 4, fourth data row vs first data row). The growth increased (up to 0.0672), but not at the level of the control model (not even after setting the upper bound to a higher value of 10, when the actual flux is lower than the set upper bound) (Table 4, data rows four and five). This means that in the treated model there are other reactions that become limiting when the maximum allowed flux through the ubiquinol:ferricytochrome c reductase is set higher. These reactions are the ATP synthase and the NADH:ubiquinone oxidoreductase, which are both operating at their maximum allowed flux in this condition (Table 5).

**Table 4:** Effect of ubiquinol:ferricytochrome c reductase reaction on growth. The first two data rows show the upper bounds set for the reaction from the RNA-Seq data for the control and treated models, respectively, as well as the resulting actual fluxes and growth rates. The other three rows show the effect in the actual flux and in growth of modifying the upper bound values. All fluxes are in mmol/(gDW*hour).

| **Id** | **Name** | **Model** | **Upper bound** | **Actual flux** | **Growth** |
| --- | --- | --- | --- | --- | --- |
| r_0439 | ubiquinol:ferricytochrome c reductase | *Control* | 3.536 | 3.465 | 0.0704 |
|  |  | *Treated* | 2.582 | 2.582 | 0.0591 |
|  |  | Control | 2.582 | 2.582 | 0.0597 |
|  |  | Treated | 3.536 | 3.536 | 0.0672 |
|  |  | Treated | 10.000 | 4.534 | 0.0688 |

Then, we keep the upper bound of ubiquinol:ferricytochrome c reductase reaction in the treated model set to 3.536 (the value from the control model, Table 4 data row one) and set the upper bounds of the other two reactions from Table 5 to an arbitrary large value (10), one at a time, to see if the control growth phenotype can be recovered (Table 6). Increasing the upper bounds of ubiquinol:ferricytochrome c reductase together with NADH:ubiquinone oxidoreductase increased to growth to 0.0672, which is still lower than the control growth rate (0.0704). But, increasing the upper bound of ubiquinol:ferricytochrome c reductase reaction together with the ATP synthase does recover the control growth phenotype, actually slightly improving the growth (0.0756 vs 0.0704) (Table 6).

**Table 5:** Potential limiting reactions in the treated model when the upper bound for the ubiquinol:ferricytochrome c reductase reaction is set to the one it has in the control model. All fluxes are in mmol/(gDW*hour).

| **Id** | **Name** | **Flux** | **Upper bound** | **Compartment** | **Model** |
| --- | --- | --- | --- | --- | --- |
| r_0226 | ATP synthase | 4.034 | 4.034 | m, c | Treated |
| r_0439 | ubiquinol:ferricytochrome c reductase | 3.536 | 3.536 | m, c | Treated |
| r_0773 | NADH:ubiquinone oxidoreductase | 0.918 | 0.918 | m | Treated |

**Table 6:** Recovering the growth phenotype in the treated model. All fluxes are in mmol/(gDW*hour).

| **Id** | **Name** | **Model** | **Upper bound** | **Actual flux** | **Growth** |
| --- | --- | --- | --- | --- | --- |
| r_0439 | ubiquinol:ferricytochrome c reductase | Treated | 3.536 | 3.536 | 0.0672 |
| r_0773 | NADH:ubiquinone oxidoreductase |  | 10.000 | 1.445 |  |
| r_0439 | ubiquinol:ferricytochrome c reductase | Treated | 3.536 | 3.536 | 0.0756 |
| r_0226 | ATP synthase |  | 10.000 | 4.863 |  |

##### 3.3.1.1 Limiting gene

The gene reaction rule for ubiquinol:ferricytochrome c reductase in the yeastGEM model is:

- (Q0105 and YBL045C and YDR529C and YEL024W and YEL039C and YFR033C and YGR183C and YHR001W__45__A and YJL166W and YOR065W and YPR191W) or (Q0105 and YBL045C and YDR529C and YEL024W and YFR033C and YGR183C and YHR001W__45__A and YJL166W and YJR048W and YOR065W and YPR191W)

This means that the protein responsible for carrying out the ubiquinol:ferricytochrome c reductase reaction is a multisubunit complex, with two possible quaternary structures, both conformed by polypeptides encoded by a set of 11 genes. The genes encoding for the components of the first quaternary structure are Q0105, YBL045C, YDR529C, YEL024W, YEL039C, YFR033C, YGR183C, YHR001W__45__A, YJL166W, YOR065W and YPR191W. The genes encoding for the second are Q0105, YBL045C, YDR529C, YEL024W, YFR033C, YGR183C, YHR001W__45__A, YJL166W, YJR048W, YOR065W and YPR191W. The maximum flux of a multisubunit complex will depend on the gene with the lowest average expression, which will be the limiting factor of the complex assembling. For both possible complex configurations, in both control and treatment conditions, **YOR065W (*CYT1*)**has the lowest average expression (Table 7). The expression level of *CYT1* is limiting the maximum flux allowed through the ubiquinol:ferricytochrome c reductase reaction in our FBA simulations. We were able to reproduce the results showed in Tables 4 – 6, by modifying *CYT1* expression values used to build the control and treated models, instead of the derived reaction upper bound.

**Table 7:** Gene average expression for components of the ubiquinol:ferricytochrome c reductase complex. The enzyme has two possible quaternary structures, labeled as complex configuration 1 and 2 in this table. The presence of the genes in a given configuration is stated in the last column.

| Gene id | Gene | Expression control | Expression treated | Complex configuration |
| --- | --- | --- | --- | --- |
| YBL045C | COR1 | 134.50 | 122.91 | 1 and 2 |
| YOR065W | *CYT1* | 122.69 | 89.59 | 1 and 2 |
| YPR191W | QCR2 | 133.78 | 138.20 | 1 and 2 |
| YFR033C | QCR6 | 125.40 | 107.32 | 1 and 2 |
| YJL166W | QCR8 | 317.91 | 552.80 | 1 and 2 |
| YEL024W | RIP1 | 160.27 | 188.18 | 1 |
| YJR048W | CYC1 | 309.61 | 179.50 | 2 |
| YDR529C | QCR7 | 407.45 | 555.42 | 2 |
| Average | All genes without expression data | 472.26 | 448.02 | 1 and 2 |

## 4 Discussion

MCHM significantly affects amino acid metabolism, increasing the total intracellular concentration of 12 out of 20 standard amino acids. As 28 genes coding for enzymes of the amino acid biosynthesis pathways are up-regulated due to MCHM treatment, the higher levels of such amino acids can be partially explained by their probable increased biosynthesis. The other contributing factor could be a reduced protein production, due to the deleterious effect of MCHM on ribosome biogenesis (down-regulating seven critical genes of the process), leading to an amino acid accumulation.

Cu^2+^ also increase the levels of some amino acids (L-glutamate, L-phenylalanine, L-leucine and L-phenylalanine) and decrease the level of L-aspartate in *S. cerevisiae* (Farrés et al., 2016)◻. From these only L-glutamate and L-aspartate variate in our analysis, both increasing their levels due to MCHM. The only amino acid which level was reduced after MCHM treatment was L-proline. In *Saccharomyces cerevisiae*, L-proline is a stress protectant, effective against oxidative stress under low pH conditions (Nugroho et al., 2016)◻, protecting against representative inhibitors like furfural, acetic acid and phenol (Wang et al., 2015)◻ and contributing to ethanol tolerance (Ohta et al., 2016)◻. It is of note that the other protective metabolite against representative inhibitors found by Wang *et al* (Wang et al., 2015)◻, myo-inositol, also has a lower level due to MCHM. It is also interesting the higher levels after MCHM treatment of glutathione, a critical protector against oxidative stress, in contrast to the previously reported oxidative stress in MCHM treated cells (Lan et al., 2015)◻. On the other hand, the level of acetyl-L-carnitine, which protects yeast cells from apoptosis and aging and inhibits mitochondrial fission (Palermo et al., 2010)◻, is increased due to MCHM. In despite of the contradictory levels of some previous metabolites, it is clear that yeast is responding to the stress caused by MCHM in our study, upregulating 11 genes related with stress response. From these genes, particularly from *AHA1*, *ENO1*, *GRE2*, *PDR3* and *PDR16*, we can infer that MCHM is affecting DNA replication (Tkach et al., 2012)◻. This is in correspondence with the previously described DNA damage effect (particularly double strand break) of MCHM in human cells (Lan et al., 2015)◻. From the other stress-related upregulated genes: *SNQ2* and *QDR3* encode multidrug transporters involved in multidrug resistance (Rogers et al., 2001; Tenreiro et al., 2005); *ENB1* encodes for an endosomal ferric enterobactin transporter, which is expressed under conditions of iron deprivation (Philpott et al., 2002)◻; *TPO1* codes for a polyamine transporter which exports spermine and spermidine from the cell during oxidative stress, controlling the timing of expression of stress-responsive genes (Krüger et al., 2013)◻; *ICT1* codes the lysophosphatidic acid acyltransferase responsible for enhanced phospholipid synthesis during organic solvent stress (Ghosh et al., 2008)◻.

We did not detect the effect of an enhanced phospholipid biosynthesis in our metabolomics analysis, by the contrary the levels of all phospholipids included in the relevant metabolites (one phosphatidylethanolamine, three phosphatidylinositol, four phosphatidylserine plus two lyso-phosphatidylethanolamine and four phosphatidic acid molecules) are decreased due to MCHM, while the levels of the remaining 37 detected phospholipids do not change. The reduced levels of these molecules of phosphatidylethanolamine, phosphatidylinositol and phosphatidylserine, together with the lower level of ergosterol in MCHM treated cells point toward a significant effect of MCHM in yeast cellular membranes, with potential effects on their biophysical properties, which could impact several cellular processes involving membranes.

In our ESI-MS experiments four branched fatty acid esters of hydroxy fatty acids (FAHFAs) were identified as relevant metabolites, which levels were increased due to MCHM. Given that FAHFAs are a relatively new discovered class of endogenous mammalian lipids with anti-diabetic and anti-inflammatory effects (Yore et al., 2014)◻, it is possible that this four FAHFAs are just false positives and unknown isobaric metabolites are the real relevant metabolites. These FAHFAS were identified with an error of no more than 1.14 dppm and a delta of 5.8e^−4^ m/z. To validate the presence of FAHFAs in yeast an additional experimental approach is required, with generation of MS/MS data and query against specialized libraries (Ma et al., 2015)◻, but this is out the scope of the current study.

The FBA simulations done with genome-scale metabolic network models (GSMNM) of MCHM treated vs non-treated control yeast were able to reproduce the deleterious effect of MCHM on cell’s growth. These GSMNM have integrated the gene expressions from the RNA-Seq data, as explained in Materials and Methods. The flux ratio through several reactions in the six significant pathways from the metabolomics analysis is linked to the simulated growth ratio in MCHM-treated vs untreated control models, but this does not indicate causality. The FBA simulations suggest a critical role to the ubiquinol:ferricytochrome c reductase as the enzyme catalyzing the limiting reaction which determine the reduced growth in MCHM. From this multisubunit complex *CYT1* product is the component limiting the overall reaction flow, and the lower expression of *CYT1* due to MCHM can explain the lower growth, at least in the FBA simulations. It is of note that the fold change of the expression levels of *CYT1* is not big enough (logFC < 2) for the gene to came out as relevant from the RNA-Seq data, but the GSMNM created are very sensitive to its levels. This highlight the extra value of RNA-Seq data integration in FBA simulations, allowing to assess the impact of gene levels in whole cell functional environment, where apparently irrelevant genes can prove to be the driven force behind observed phenotypes. Transcription of *CYT1* is positively controlled by oxygen in the presence of glucose, through the haem signal and mediated by the Hap1. It is additionally regulated by the HAP2/3/4 complex which mediates gene activation mainly under glucose-free conditions. Basal transcription is partially effected by Cpf1, a centromere and promoter-binding factor (Oechsner et al., 1992)◻.

The other significant reaction that comes from the FBA analysis is the ATP synthase, which maximum allowed flux or upper bound is required to be increased together with the one of ubiquinol:ferricytochrome c reductase to rescue the control growth phenotype in the MCHM treated model. Combining flux balance analysis with *in vitro* measured enzyme specific activities it was determined that fermentation is more catalytically efficient than respiration (Nilsson and Nielsen, 2016)◻, producing more ATP per mass of required enzymes. In that study the enzyme F1F0-ATP synthase was found to have flux control over respiration in the model, causing the Crabtree Effect (Nilsson and Nielsen, 2016)◻.

## 5 Conclusions

MCHM produce amino acid accumulation in *S. cerevisiae*, affecting several amino acid related metabolic pathways and probably slowing down protein biosynthesis due to the downregulation of genes related to ribosome biogenesis. MCHM affects phospholipid biosynthesis, reducing the levels of different molecules of phosphatidylethanolamine, phosphatidylinositol and phosphatidylserine, which, in addition to lower levels of ergosterol, should affect cellular membranes composition and their biophysical properties. The FBA simulations suggest that the lower flow through ubiquinol:ferricytochrome c reductase reaction, caused by the MCHM-provoked under-expression of *CYT1* gene, could be the driven force behind the observed effect on yeast metabolism and growth.

## 6 Acknowledgments

We would like to thank BioNano Research Facilities staff at West Virginia University, Dr. Callie Walsh and Sandra Majuta for their support with the ESI-MS experiments.

This work was funded by a grant from the NIH (R15ES026811-01A1) to JEGG. The funding source was not involved in the study design; nor in the collection, analysis and interpretation of data; nor in the writing of the report; and neither in the decision to submit the article for publication.

## 7 Declarations of interest

None.

## 8 Author contributions

AP and JEGG designed the experiments. AP performed the extractions. AP and KMK carried out the GC/MS experiments. AP did the ESI-MS experiments and the FAB simulations. JEGG did the RNA-Seq experiment. AP analyzed all the data and wrote the manuscript. JEGG and KMK reviewed the manuscript.

## Abbreviations

MCHM: 4-Methylcyclohexanemethanol
GSMNM: genome-scale metabolic network models
FBA: flux balance analysis

