## Supplementary figures and images for "Effects of MCHM on yeast metabolism"

### Supplemental Figure 1

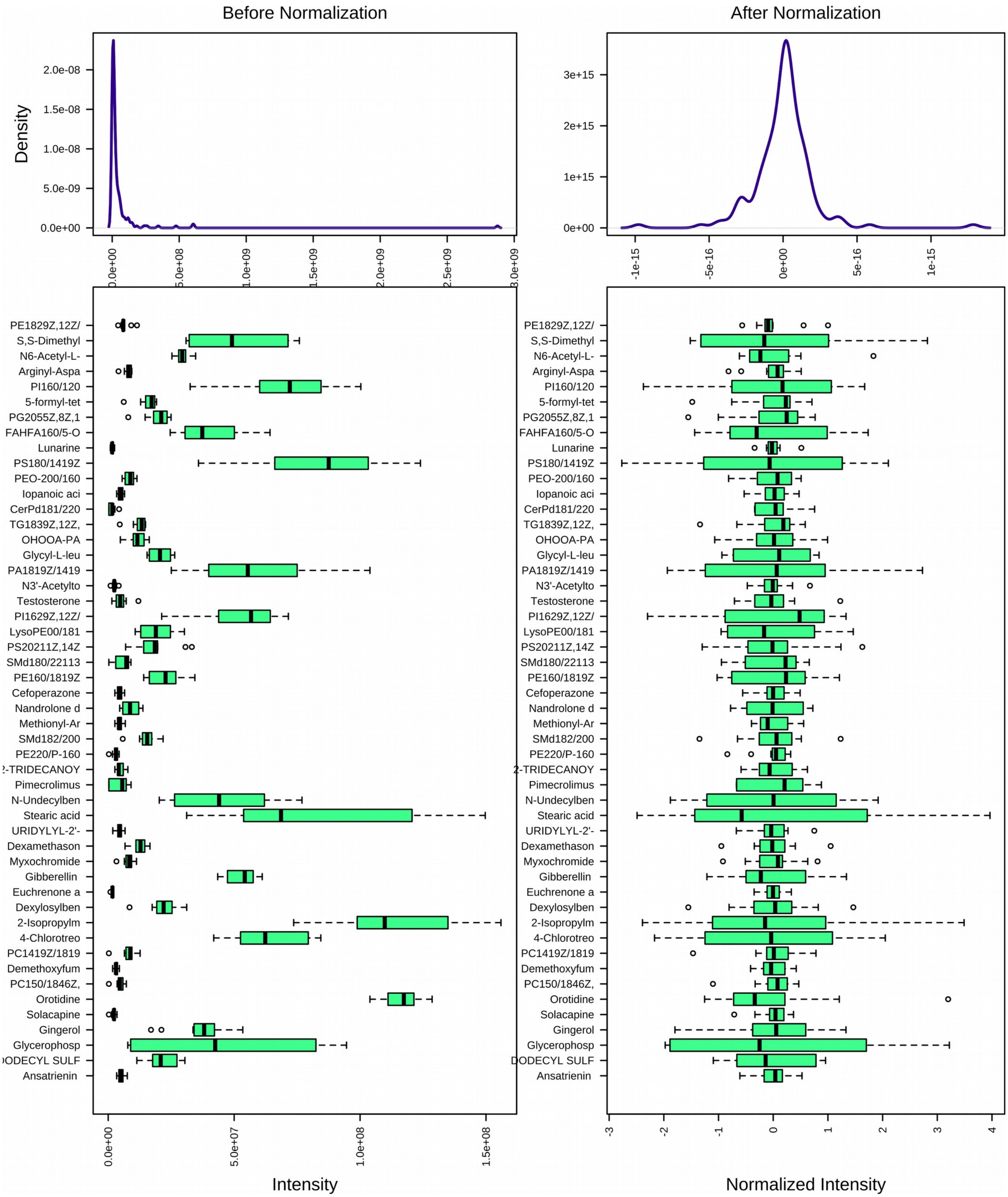


*Supplemental Figure 1: Peaks intensities of ESI-MS metabolites before and after normalization.*

### Supplemental Figure 2

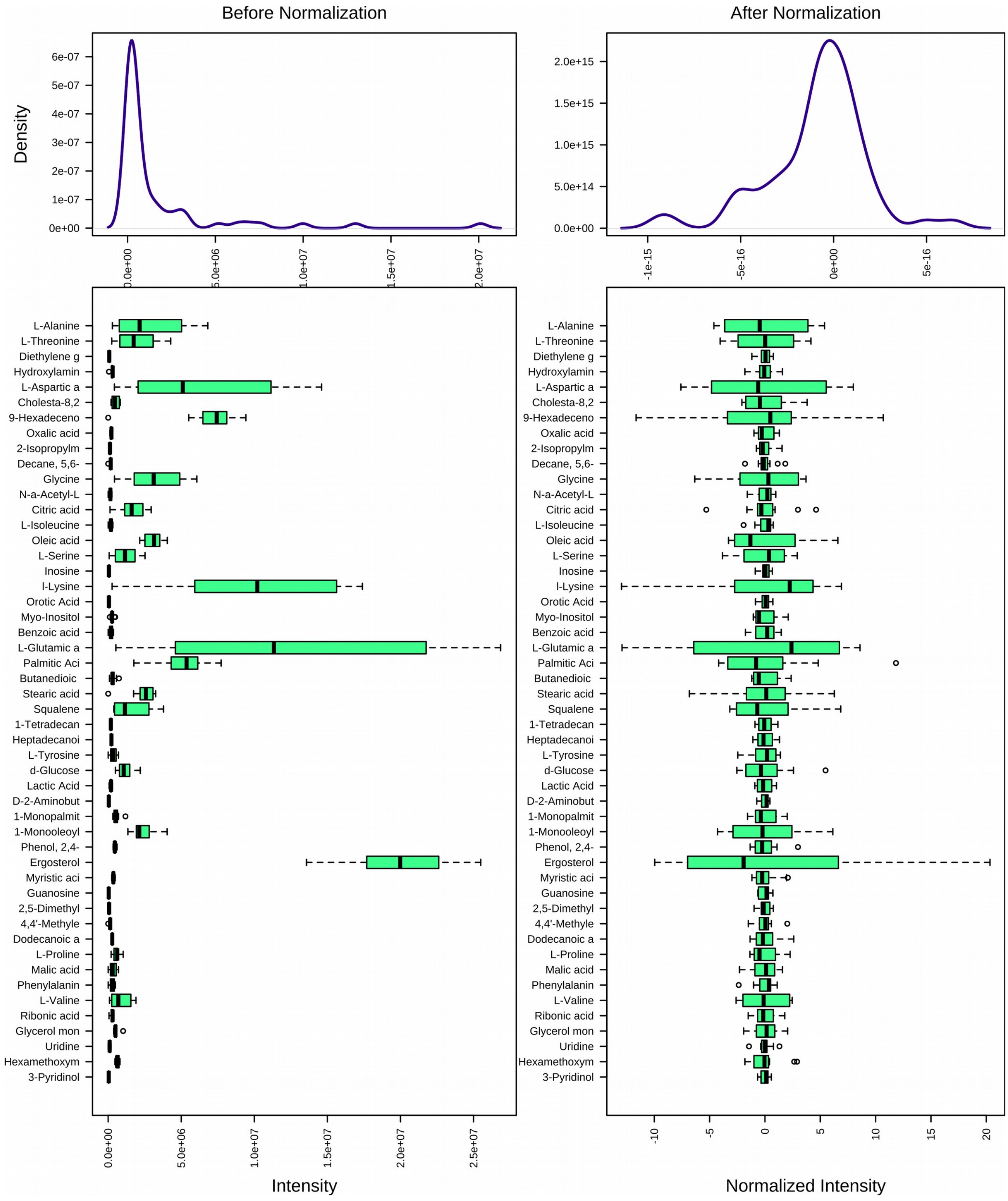


*Supplemental Figure 2: Peaks intensities of GC-MS metabolites before and after normalization.*
