## Supplemental Figure 3 for "Effects of MCHM on yeast metabolism"

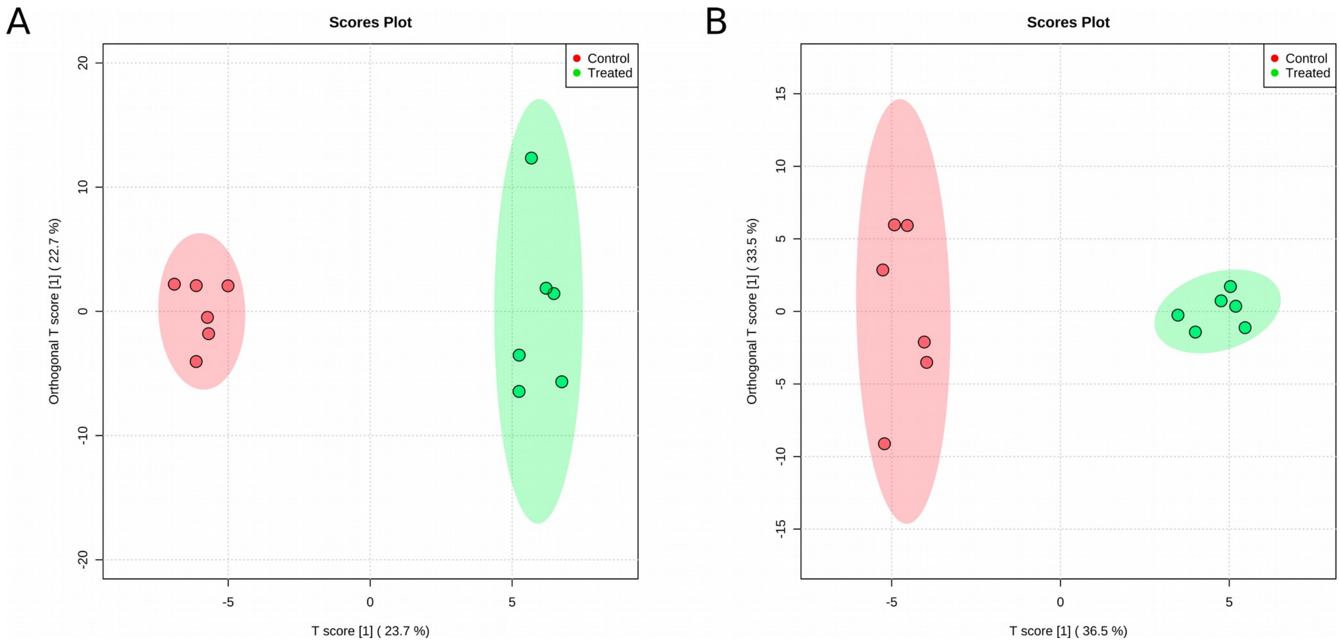


*Supplemental Figure 3: Orthogonal-Orthogonal Projections to Latent Structures Discriminant Analysis (OPLS-DA) of ESI-MS (A) and GC-MS (B) data. The 95 % confidence area are shown as well as the explained variance, shown in brackets in the corresponding axis labels.*
