## Supplemental Figure 4 for "Effects of MCHM on yeast metabolism"

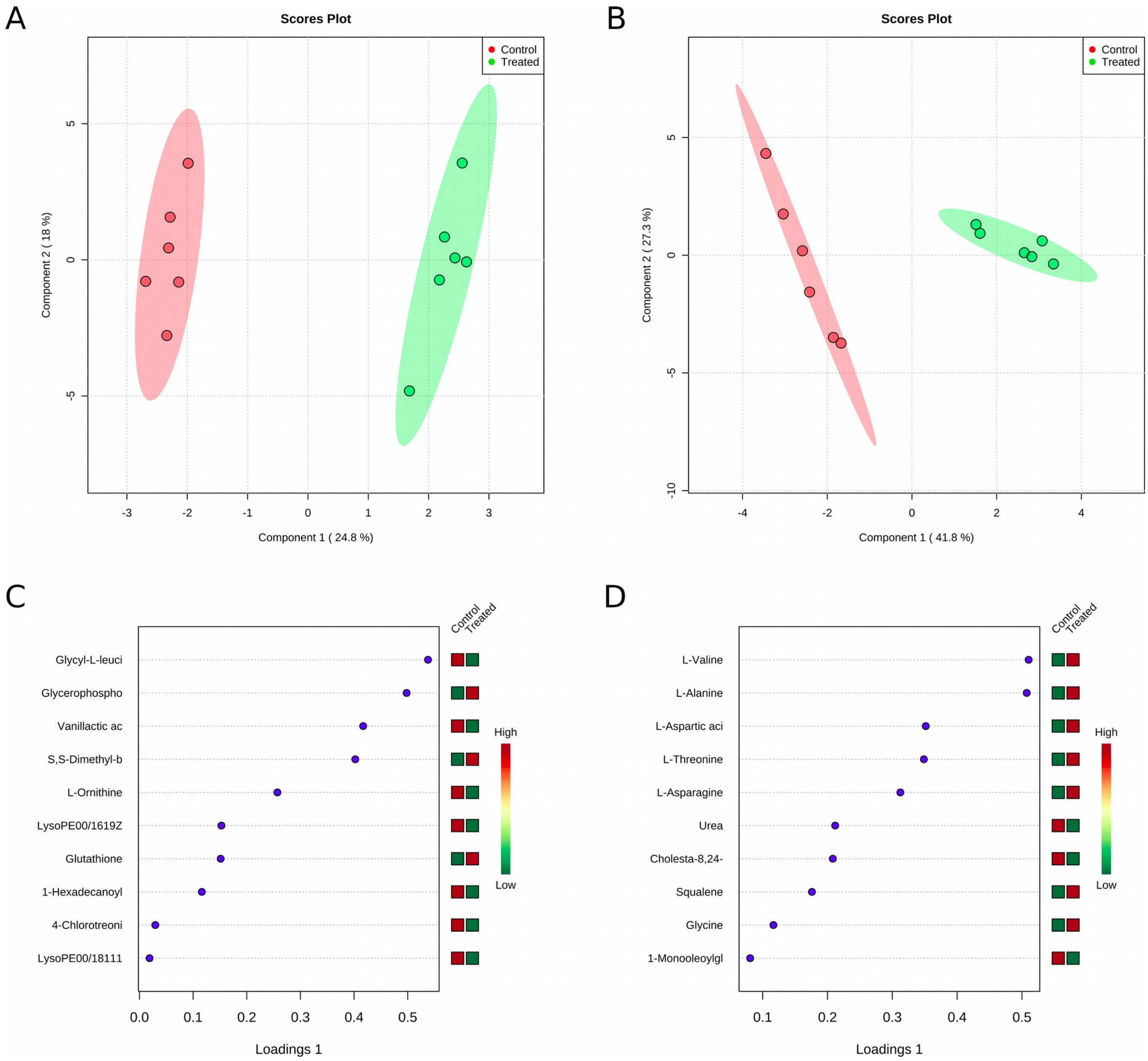


*Supplemental Figure 4: Sparse Partial Least Squares - Discriminant Analysis (sPLS-DA). Scores plot between the first two components for A) ESI-MS and B) GC-MS respectively, with the 95 % confidence area shown and the explained variance shown in brackets in the corresponding axis labels. Plot showing the variables selected by the sPLS-DA model for a given component in C) ESI-MS and D) GC-MS experiments. The variables are ranked by the absolute values of their loadings.*
