## Supplemental Figure 5 for "Effects of MCHM on yeast metabolism"

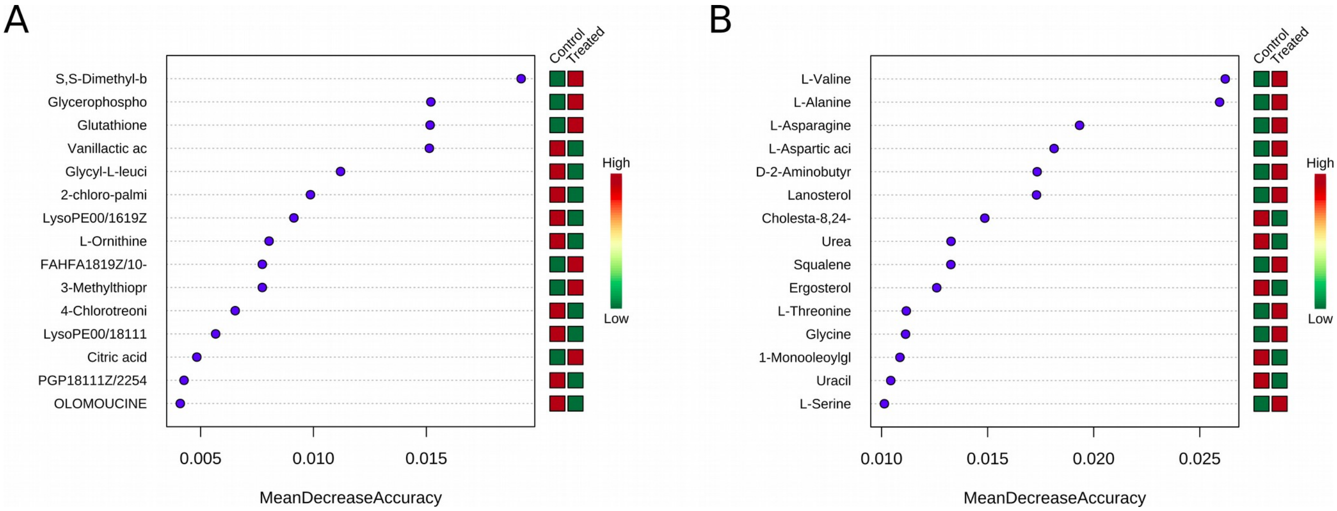


*Supplemental Figure 5: Significant features identified by Random Forest for A) ESI-MS and B) GC-MS data. The features are ranked by the mean decrease in classification accuracy when they are permuted.*
