## Supplemental Figure 6 for "Effects of MCHM on yeast metabolism"

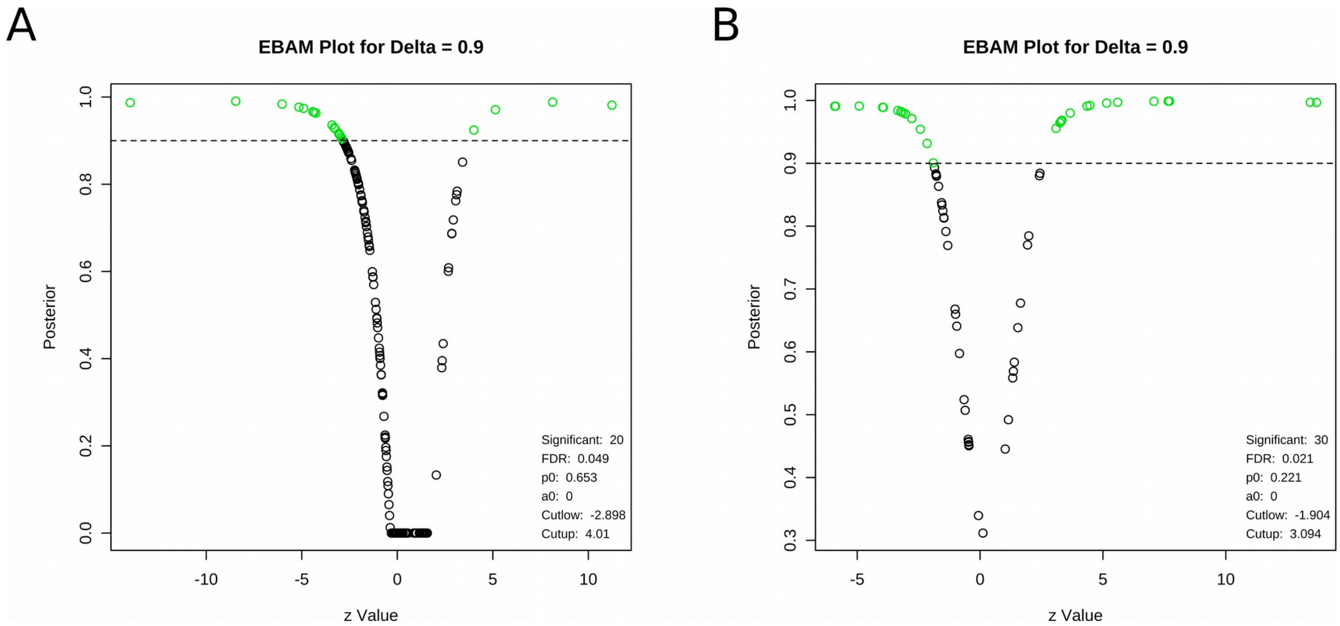


*Supplemental Figure 6: Empirical Bayesian Analysis of Microarray (EBAM) for A) ESI-MS and B) GC-MS data. 20 and 30 significant compounds are identified with this method for ESI-MS and GC-MS, respectively.*
