## Supplemental Figure 7 for "Effects of MCHM on yeast metabolism"

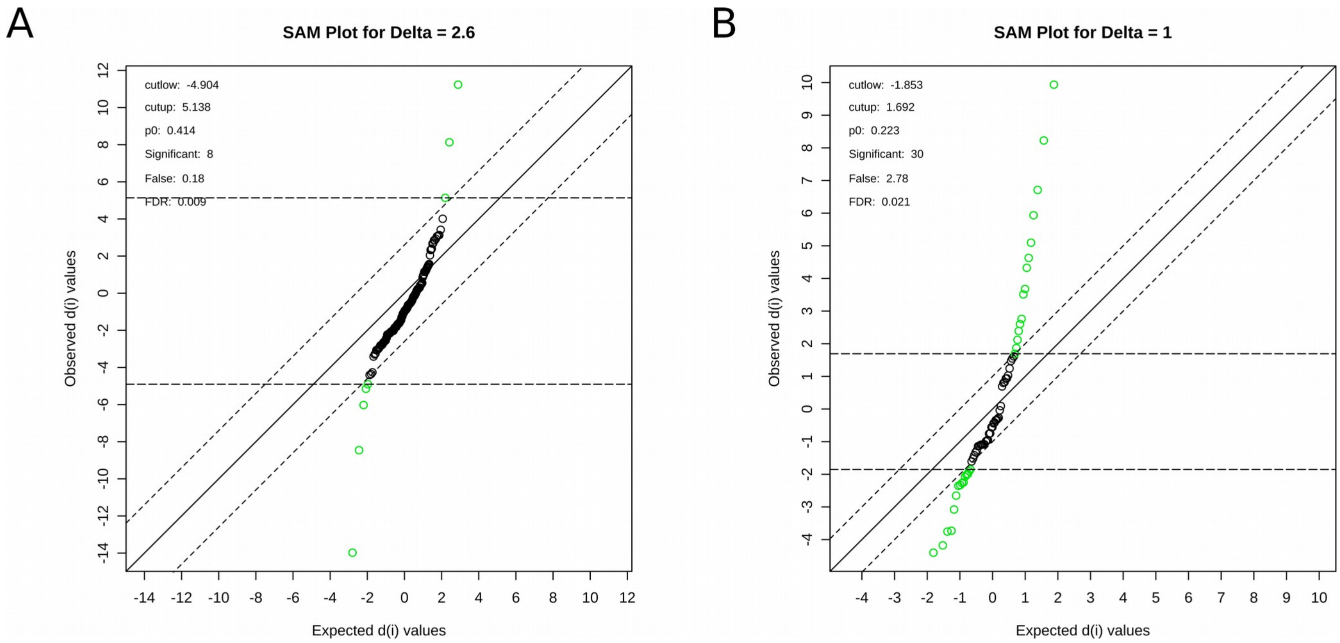


*Supplemental Figure 7: Significance Analysis of Microarray (SAM) for A) ESI-MS and B) GC-MS data. The green circles represent features that exceed the specified threshold. 8 and 30 significant features are identified by SAM from ESI-MS and GC-MS respectively.*
