## Supplemental Figure 9 for "Effects of MCHM on yeast metabolism"

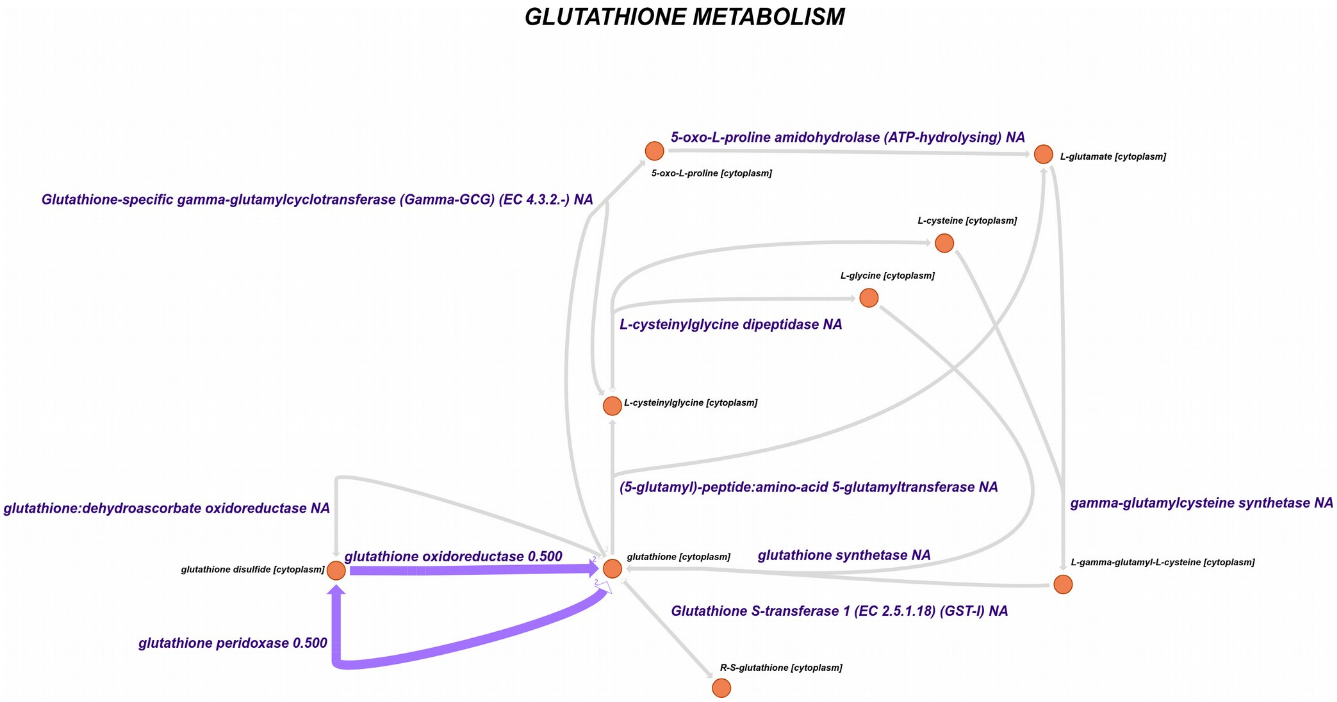


*Supplemental Figure 9: Escher map of the glutathione metabolism, with the flux ratios between treated and control model FBA solutions represented. The thick and color of the edges are a function of the respective ratio values. There is only data for the inter-conversion between glutathione and glutathione disulfide, which fluxes are reduced to half in the treated model.*
