## Supplemental Figure 10 for "Effects of MCHM on yeast metabolism"

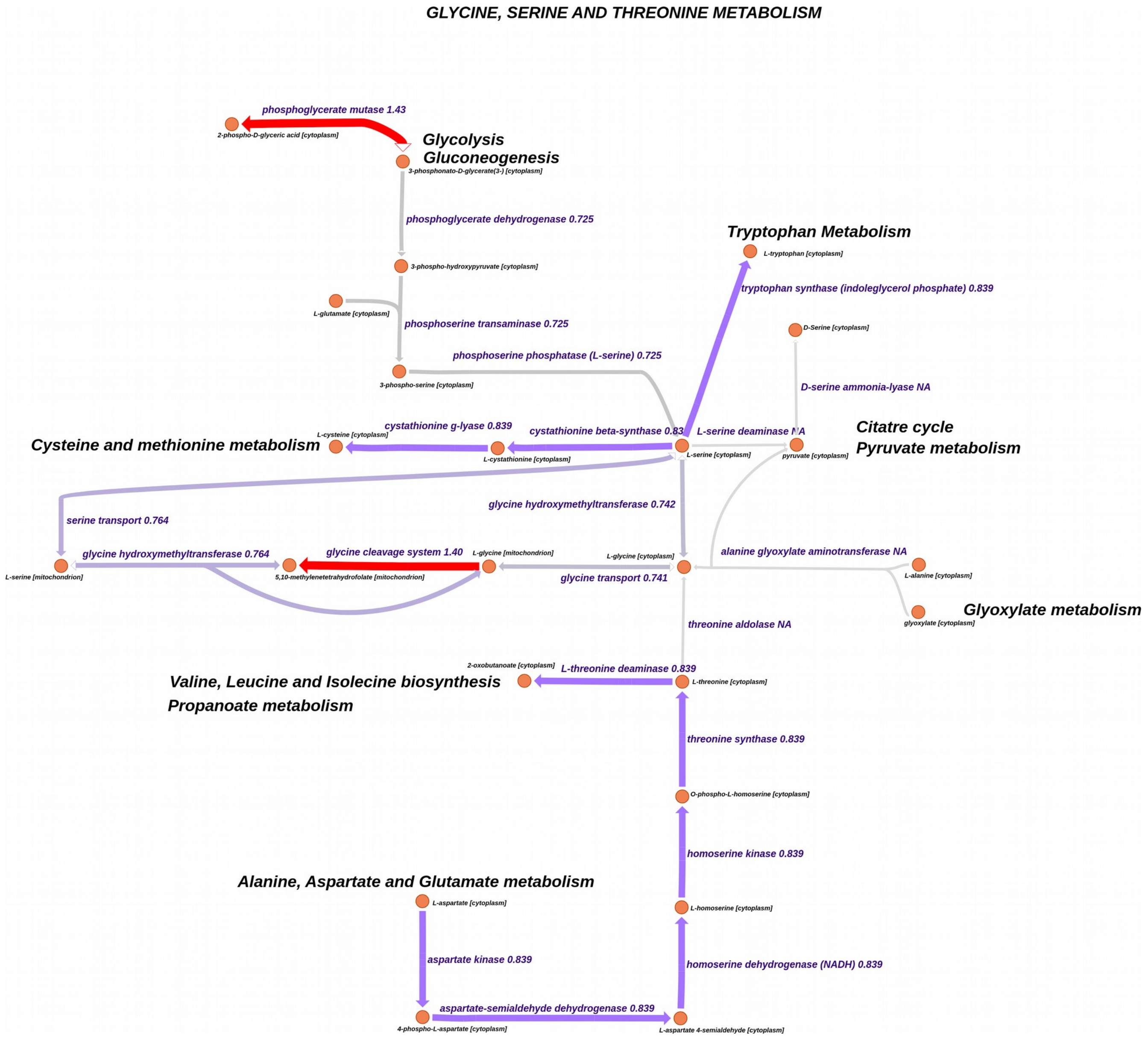


*Supplemental Figure 10: Escher map of the Glycine, Serine and Threonine metabolism, with the flux ratios between treated and control model FBA solutions represented. The thick and color of the edges are a function of the respective ratio values. The ratio value of 0.839 is common among the map.*
