## Supplemental Figure 11 for "Effects of MCHM on yeast metabolism"

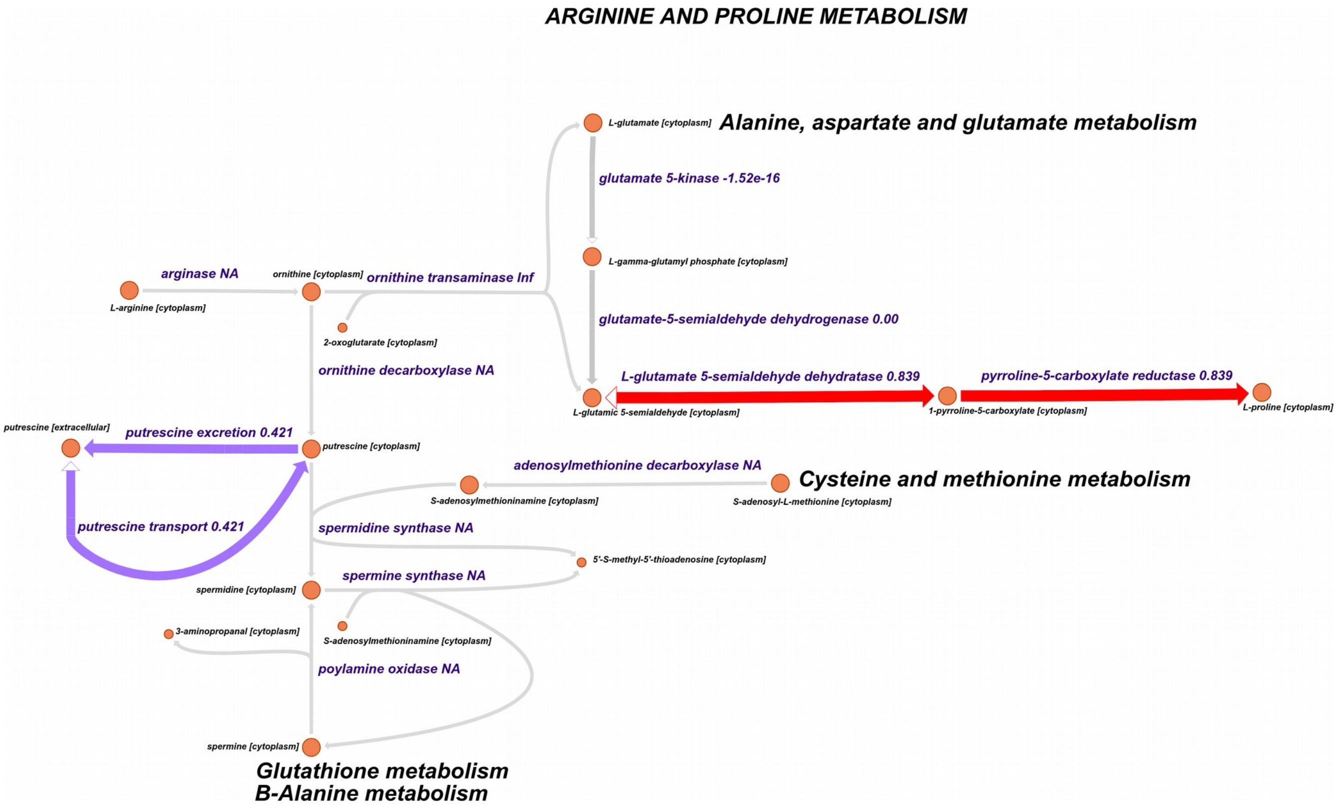


*Supplemental Figure 11: Escher map of the Arginine and Proline metabolism, with the flux ratios between treated and control model FBA solutions represented. The thick and color of the edges are a function of the respective ratio values.*
