## Supplemental Figure 12 for "Effects of MCHM on yeast metabolism"

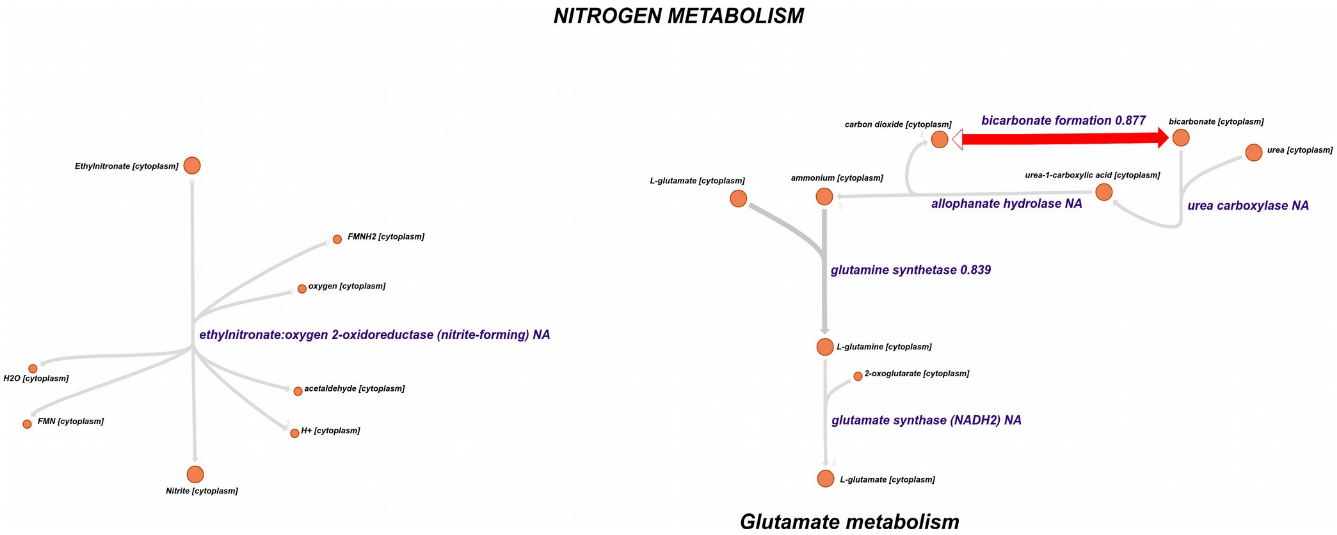


*Supplemental Figure 12: Escher map of the nitrogen metabolism, with the flux ratios between treated and control model FBA solutions represented. The thick and color of the edges are a function of the respective ratio values.*
