## Supplemental Figure 13 for "Effects of MCHM on yeast metabolism"

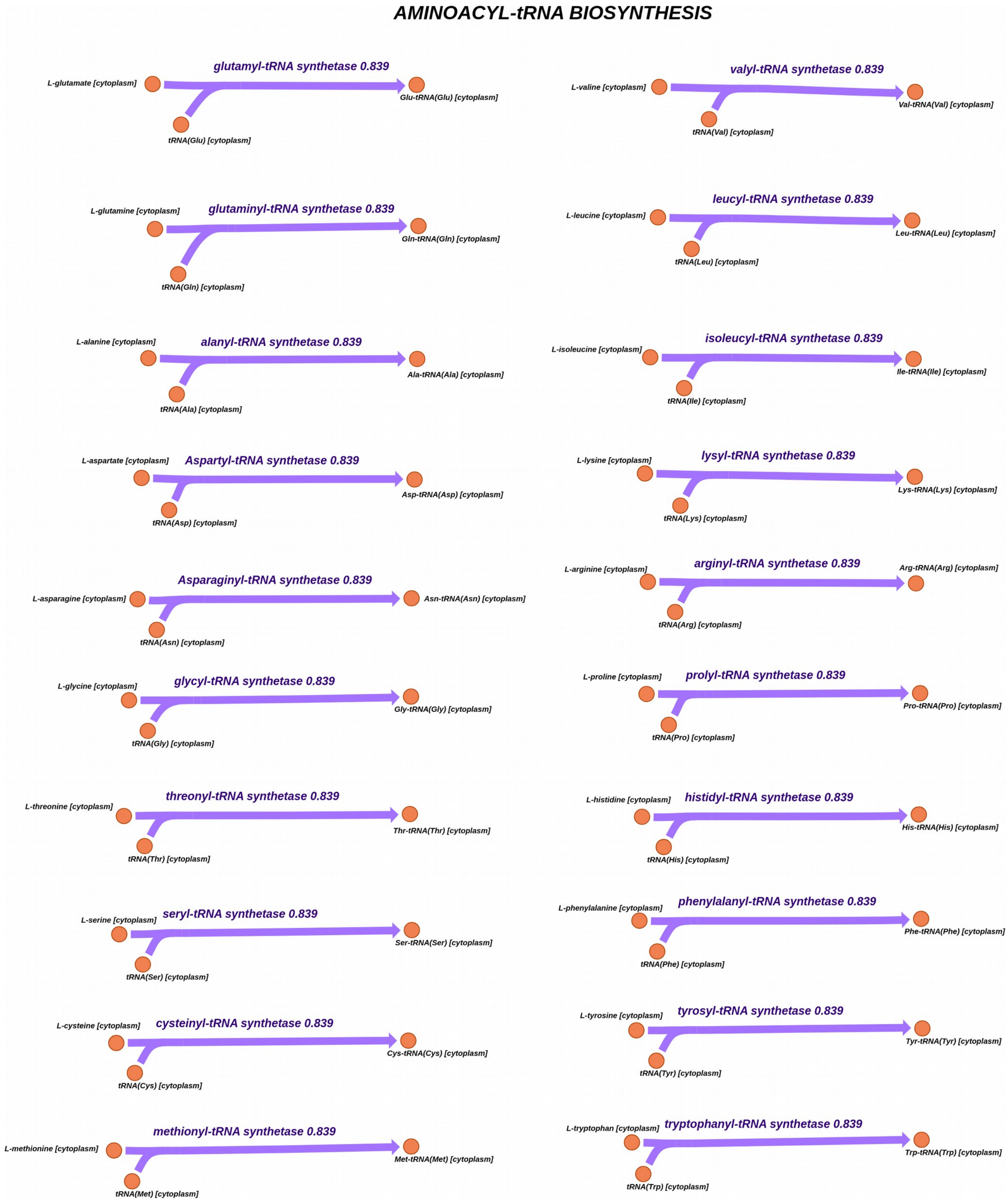


*Supplemental Figure 13: Escher map of the Aminoacil t-RNA biosynthesis, with the flux ratios between treated and control model FBA solutions represented. The thick and color of the edges are a function of the respective ratio values. All the reactions have the ratio value of 0.839.*
